# Reinforcement Learning and Bayesian Inference Provide Complementary Models for the Unique Advantage of Adolescents in Stochastic Reversal

**DOI:** 10.1101/2020.07.04.187971

**Authors:** Maria K. Eckstein, Sarah L. Master, Ronald E. Dahl, Linda Wilbrecht, Anne G.E. Collins

## Abstract

During adolescence, youth venture out, explore the wider world, and are challenged to learn how to navigate novel and uncertain environments. We investigated whether adolescents are uniquely adapted to this transition, compared to younger children and adults. In a stochastic, volatile reversal-learning task with a sample of 291 participants aged 8-30, we found that adolescents outperformed both younger and older participants. We developed two independent cognitive models, based on Reinforcement learning (RL) and Bayesian inference (BI). The RL parameter for learning from negative outcomes and the BI parameters specifying participants’ mental models peaked closest to optimal in adolescents, suggesting a central role in adolescent cognitive processing. By contrast, persistence and noise parameters improved monotonously with age. We distilled the insights of RL and BI using principal component analysis and found that three shared components interacted to form the adolescent performance peak: adult-like behavioral quality, child-like time scales, and developmentally-unique processing of positive feedback. This research highlights adolescence as a neurodevelopmental window that may be specifically adapted for volatile and uncertain environments. It also shows how detailed insights can be gleaned by using cognitive models in new ways.

## 1. Introduction

In mammals and other species with parental care, there is typically an adolescent stage of development in which the young are no longer supported by parental care, but are not yet adult (Natterson-Horowitz and Bowers, 2019). This adolescent period is increasingly viewed as a critical epoch in which organisms explore the world, make pivotal decisions with short- and long-term impact on survival (Frankenhuis and Walasek, 2020), and learn about important features of their environment (DePasque and Galván, 2017; Steinberg, 2005), likely taking advantage of a second window of brain plasticity (Larsen and Luna, 2018; Lourenco and Casey, 2013; Piekarski, Johnson, et al., 2017).

In humans, adolescence often involves an expansion of environmental contexts and increasingly frequent transitions between them (*contextual volatility*; e.g., new pastime activities, growing relevance of peer relationships; Albert et al., 2013; Somerville et al., 2017), as well as increased exposure to uncertainty (*outcome stochasticity*; e.g., increased risk-taking and sensation seeking, increased unpredictability of social interactions; Romer and Hennessy, 2007; van den Bos and Hertwig, 2017). Accordingly, it has been argued that adolescent brains and minds are specifically adapted to contextual volatility and outcome stochasticity, showing an increased ability to learn from and succeed in these situations (Dahl et al., 2018; Davidow et al., 2016; Johnson and Wilbrecht, 2011; Lloyd et al., 2020; Lourenco and Casey, 2013; Sercombe, 2014).

The goal of this study was to test this *U-shape* hypothesis in a controlled laboratory environment. We employed a stochastic reversal-learning task in a large developmental sample (*n* = 291) with a wide, continuous age range (8-30 years), offering enough statistical power to observe non-linear effects of age (such as the predicted U-shaped pattern with peak in adolescence). Another goal was to identify a computational explanation of the non-linear development of underlying cognitive processes, using state-of-the-art computational modeling.

### 1.1. U-Shapes in Development

The predicted U-shaped development is in line with recent findings: Adolescents show non-linear developments both in terms of neural maturation and with regard to behaviour, including emotional processing, learning, and decision making (for reviews, see Dahl et al., 2018; Giedd et al., 1999; Somerville and Casey, 2010; Sowell et al., 2003; Toga et al., 2006). Research on adolescent development has often focused on aspects with negative reallife outcomes, including elevated risk-taking and sensation seeking (Braams et al., 2015; Galvan et al., 2006; Harden and Tucker-Drob, 2011; Romer and Hennessy, 2007), but positive aspects have become evident more recently, too (DePasque and Galván, 2017; Sercombe, 2014). For example, adolescents have outperformed adults in certain measures of creativity (Kleibeuker et al., 2013) and showed enhanced social learning (Brandner et al., 2021; Gopnik et al., 2017) and exploration (Somerville et al., 2017). With particular interest to our hypothesis, adolescents have outperformed adults on stochastic learning tasks (Cauffman et al., 2010; Davidow et al., 2016) and some aspects of a reversal-learning task (van der Schaaf et al., 2011; Fig. 3). Adolescents’ behavioral advantages on these tasks are likely related to non-linear patterns of brain development (Dahl et al., 2018; Giedd et al., 1999; Somerville and Casey, 2010; Sowell et al., 2003; Toga et al., 2006), and potentially modulated by puberty-related hormonal changes (Blakemore et al., 2010; Braams et al., 2015; Gracia-Tabuenca et al., 2021; Laube, Lorenz, et al., 2020; Op de Macks et al., 2016; Piekarski, Johnson, et al., 2017), some of which have been associated with cognitive flexibility, decision making under uncertainty, and feedback processing, cognitive processes that are particularly relevant for stochastic reversal learning. Supporting this perspective, similar prowess in flexibility has been reported in developing rodents (Guskjolen et al., 2017; Johnson and Wilbrecht, 2011; Simon et al., 2013), and linked to neural and hormonal maturation (Delevich et al., 2019; Piekarski, Boivin, et al., 2017).

### 1.2. Stochastic Reversal Learning

Studied since the birth of the cognitive neurosciences, reversal learning has recently seen an exponential growth in published studies. Originally meant to measure response inhibition, reversal paradigms are now agreed to primarily measure cognitive flexibility (Izquierdo et al., 2017). In stochastic reversal-learning tasks, participants need to discriminate which outcomes occur due to inherent stochasticity, in which case they should double down on their current, appropriate strategy; and which outcomes are caused by context switches, in which case they need to rapidly change their strategy. Stochastic reversal tasks therefore pose a fundamental tension between persistence with previous strategies and adaptability to new circumstances, a major challenge in the adolescent transition.

An abundance of studies has mapped the specific brain areas (most notably orbitofrontal cortex and striatum) and endocrine systems (mainly serotonin, dopamine, and glutamate) relevant for reversal learning (Clark et al., 2004; Frank and Claus, 2006; Hamilton and Brigman, 2015; Izquierdo et al., 2017; Izquierdo and Jentsch, 2012; Kehagia et al., 2010; Yaple and Yu, 2019). Most of these systems still undergo developmental changes during adolescence and early adulthood, oftentimes following U-shaped trajectories (Albert et al., 2013; Casey et al., 2008; Dahl et al., 2018; DePasque and Galván, 2017; Larsen and Luna, 2018; Laube, Lorenz, et al., 2020; Lourenco and Casey, 2013; Piekarski, Johnson, et al., 2017; Somerville and Casey, 2010; Toga et al., 2006). This suggests that behavioral development, as well, might show a non-linear development.

However, even though reversal tasks have been used abundantly in developmental populations (e.g., Adleman et al., 2011; DePasque and Galván, 2017; Dickstein, Finger, Brotman, et al., 2010; Dickstein, Finger, Skup, et al., 2010; Finger et al., 2008; Harms et al., 2018; Hildebrandt et al., 2018; Minto de Sousa et al., 2015), we still know surprisingly little about their developmental trajectory. To our knowledge, only three studies have assessed this: Two employed binary group designs comparing adolescents to adults, but did not show significant age differences in performance (Hauser et al., 2015; Javadi et al., 2014). Note that the U-shaped developments we predict would be undetectable in most binary group designs. A third study employed a deterministic reversal task, and tested four age groups across adolescence, which allowed to assess non-linear changes (van der Schaaf et al., 2011). Indeed, there was an adolescent peak in reversal performance (Fig. 3). Here, we seek to extend this finding by studying a larger sample, employing a stochastic task, and to provide insights into the cognitive mechanisms that support adolescents’ superior performance, using computational modeling.

### 1.3. Computational Modeling

#### 1.3.1. Reinforcement Learning (RL)

RL is a popular framework to model probabilistic reversal learning (Boehme et al., 2017; Chase et al., 2010; Gläscher et al., 2009; Hauser et al., 2015; Javadi et al., 2014; Metha et al., 2020; Peterson et al., 2009). RL agents choose actions based on action *values* that reflect actions’ expected long-term cumulative reward. Action values are typically estimated by incrementally updating them every time an action outcome is observed (see section 4.5.1). The size of each update, determined by an agent’s *learning rate*, captures the integration time scale, i.e., whether value estimates are based on few recent outcomes, or many outcomes that reach further into the past. A specialized network of brain regions, including the striatum and frontal cortex, has been associated with specific RL-like computations (Frank and Claus, 2006; D. Lee et al., 2012; Niv, 2009; O’Doherty et al., 2015).

As a computational model, RL interprets cognitive processing during reversal learning as *value learning*: RL agents continuously adjust current action values based on new outcomes, striving to learn increasingly accurate values (Fig. 3A, left). Importantly, the same gradual learning process occurs during stable task periods and after context switches, without an explicit concept of switching. Behavioral switching occurs when the previously-rewarding action has accumulated enough negative outcomes to push its value below the previously-unrewarding action, in stark contrast to the quick and flexible switching behavior observed in humans and non-human animals (Costa et al., 2015; Izquierdo et al., 2017).

Because basic RL algorithms hence behave sub-optimally in volatile environments (Gershman and Uchida, 2019; Sutton and Barto, 2017), we implemented model augmentations that alleviate these issues, including distinct learning rates for positive and negative outcomes (e.g., Caźe and van der Meer, 2013; Christakou et al., 2013; Dabney et al., 2020; Frank et al., 2004; Harada, 2020; Javadi et al., 2014; Lefebvre et al., 2017; Palminteri et al., 2016; van den Bos et al., 2012), counter-factual updating (e.g., Boehme et al., 2017; Boorman et al., 2011; Gläscher et al., 2009; Hauser et al., 2014; Palminteri et al., 2016), and choice persistence (e.g., Sugawara and Katahira, 2021). See section 4.5.1 for details.

#### 1.3.2. Bayesian Inference (BI)

Many have also argued that a different computational framework, BI (specifically, Hidden Markov Models), provides a better model for human and animal behavior in reversal tasks (Bromberg-Martin et al., 2010; Costa et al., 2015; Fuhs and Touretzky, 2007; Gershman and Uchida, 2019; Solway and Botvinick, 2012). Indeed, BI models have shown better fit than RL models in three empirical studies on human adults (Hauser et al., 2014; Schlagenhauf et al., 2014) and macaques (Bartolo and Averbeck, 2020). Furthermore, BI is the standard modeling framework in the “inductive reasoning” literature, whose tasks are sometimes identical to stochastic reversal-learning tasks (e.g., Nassar et al., 2012; O’Reilly et al., 2013; Yu and Dayan, 2005).

The main reason for the supposed superiority of BI in reversal learning is the ability to reason about *hidden states* and switch behavior rapidly after recognizing state changes. Hidden states are unobservable features that determine an environment’s underlying mechanics (e.g., in reversal tasks, which choices are objectively correct and incorrect). These states can be difficult to infer when they lead to observable outcomes probabilistically. BI agents infer hidden states by engaging *predictive models* that determine how likely different outcomes occur in each state (e.g., how likely a negative outcome occurs after a correct versus incorrect choice). Agents continuously combine state *likelihoods* with their *prior beliefs* about hidden states to obtain updated *posterior beliefs* (Perfors et al., 2011; Sarkka, 2013).

Even though the BI framework supposedly provides an excellent choice to model stochastic reversal learning, it is still used rarely, and—to our knowledge—never in a developing population. Hence, BI could provide insights into the development of reversal learning that have so far escaped our attention, for example characterizing predictive mental models and inferential reasoning.

The goal of this study was to characterize adolescent behavior in stochastic reversal, and to identify its underlying cognitive mechanisms, using computational modeling: Whereas RL can tell us about participants’ learning rates in different situations and is in line with previous developmental modeling work (Hauser et al., 2015; Javadi et al., 2014), the majority of non-developmental work on reversal learning, and most standard cognitive neuroscience tasks, BI can assess participants’ mental task models and inferential processes, and is increasingly seen as a superior model compared to RL for reversal paradigms. Using in-depth modeling analyses, we found that the insights of both models could be combined to identify features of cognitive processing that went beyond any specific model, including time scales and feedback processing. Our results support the existence of an adolescent performance peak in stochastic reversal learning, and show that it stems from the parallel development of multiple cognitive mechanisms.

## 2. Results

### 2.1. Task Design

Participants’ goal in the experimental task was to collect gold coins, which were hidden in one of two locations (Fig. 1A). Which location contained the coin could change unpredictably (*volatility*), and the correct location did not always provide coins (*stochasticity*). On each trial, two identical boxes appeared on the screen. Participants chose one, either receiving a coin (reward) or not (Fig. 1A). The correct location was rewarded in 75% of the trials on which it was chosen, whereas the other one was never rewarded. Positive outcomes were therefore diagnostic of correct actions, whereas negative outcomes were ambiguous, arising from either stochastic noise or task switches. After reaching a non-deterministic performance criterion (see section 4.3), an unsignaled switch occurred, and the opposite location became rewarding (5-9 switches; 120 trials (Fig. 1B). Before the main task, participants completed a child-friendly tutorial (section 4.3).

**Figure 1:**
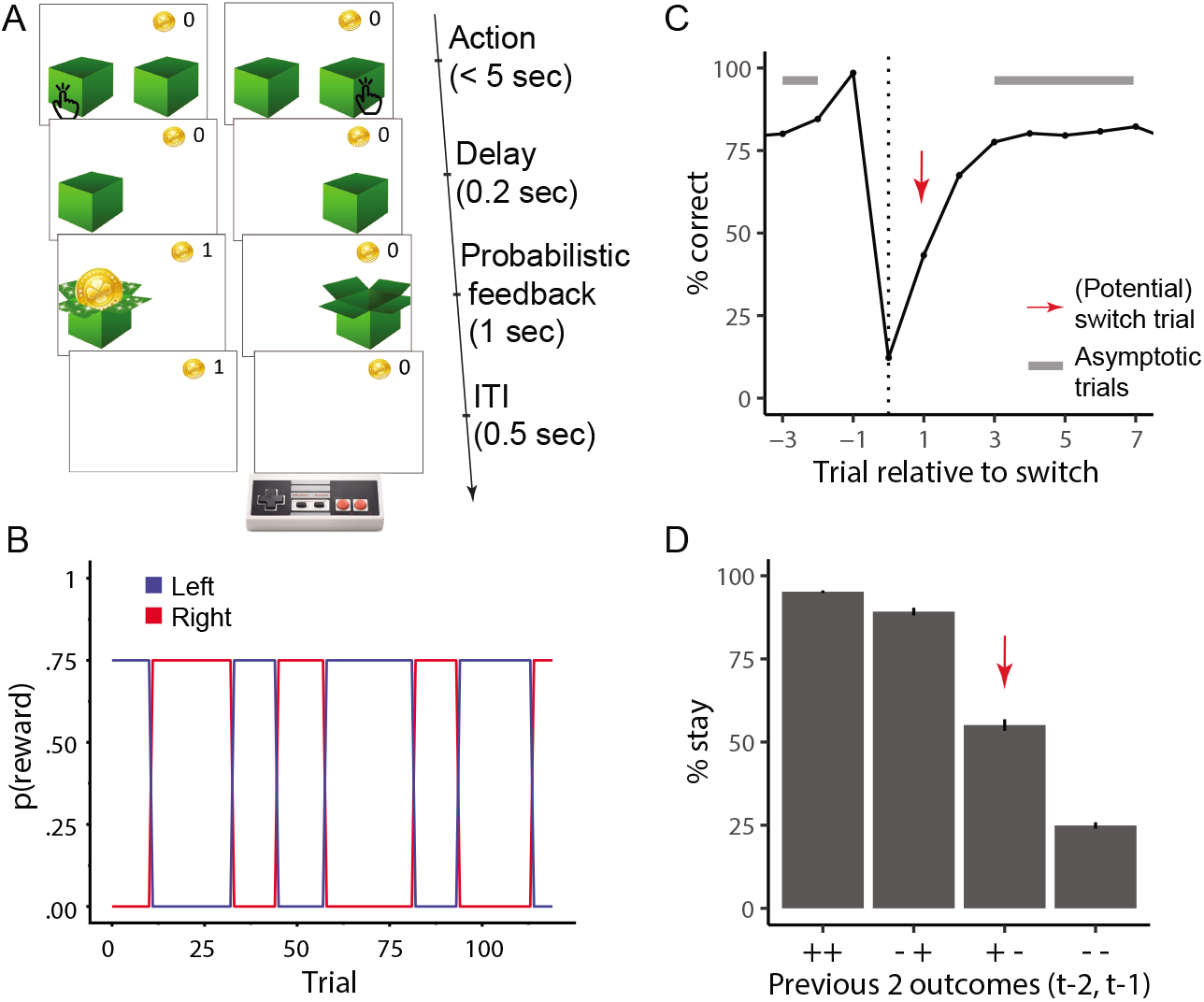
(A) Task design. On each trial, participants chose one of two boxes, using the two red buttons of the shown game controller. The chosen box either revealed a gold coin (left) or was empty (right). The probability of coin reward was 75% on the rewarded side, and 0% on the non-rewarded side. (B) The rewarded side changed multiple times, according to unpredictable task switches. (C) Average human performance and standard errors, aligned to true task switches (dotted line; trial 0). Switches only occurred after rewarded trials (section 4.3), resulting in performance of 100% on trial −1. The red arrow shows the switch trial, grey bars show trials included as asymptotic performance. (D) Average probability of repeating a previous choice (“stay”) as a function of the two previous outcomes (*t* 2, *t* 1) for this choice (“+”: reward; “-”: no reward). Error bars indicate between-participant standard errors. Red arrow highlights potential switch trials, i.e., when a rewarded trial is followed by a non-rewarded one, which—from participants’ perspective—is consistent with a task switch.

### 2.2. Task Behavior

Participants gradually adjusted their behavior after task switches, and on average started selecting the correct action about 2 trials after a switch, reaching asymptotic performance of around 80% correct choices within 3-4 trials after a switch (Fig. 1C). Participants almost always repeated their choice (“stayed”) after receiving positive outcomes (“-+” and “+ +”), and often switched actions after receiving two negative outcomes (“--”). Behavior was most ambivalent after receiving a positive followed by a negative outcome (“+ -”), i.e., on “potential” switch trials (Fig. 1D; for age differences, see suppl. Fig. 15).

#### 2.2.1. Age Differences: Performance Peak in Adolescents

Using (logistic) mixed-effects regression to test the continuous effects of age on performance (for detailed methods, see section 4.4), we found positive linear and negative quadratic age contrasts in all three performance measures (overall accuracy, stay after potential switch, accuracy on asymptotic trials; Table 1). This is in accordance with a general increase in performance from childhood to adulthood that is modified by an adolescent peak in performance.

**Table 1:**
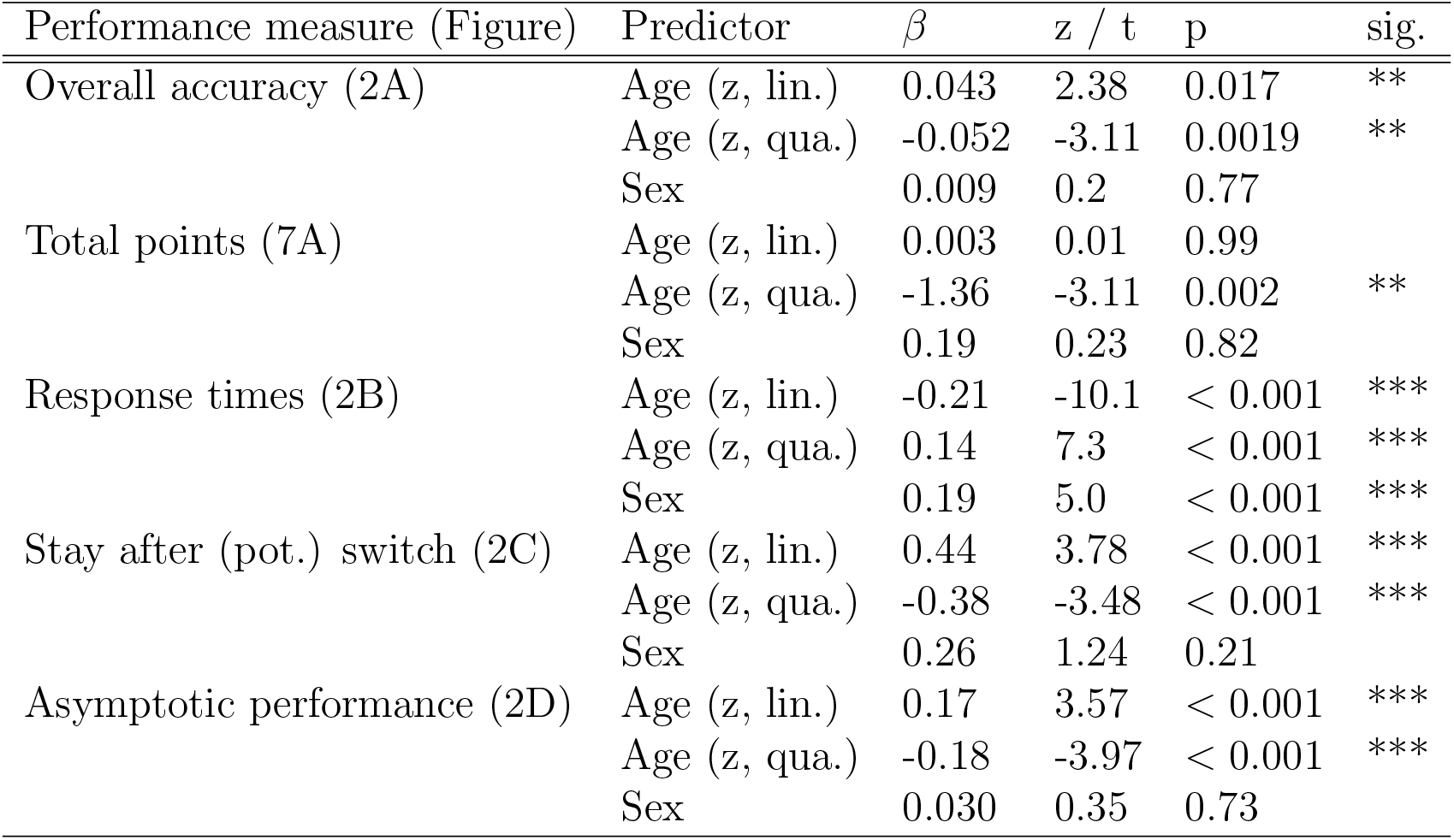
Statistics of mixed-effects regression models predicting performance measures from sex (male, female), age (z-scored; “lin.”), and quadratic age (square of z-scored age; “qua.”; for details, see section 4.4). Overall accuracy, stay after potential (pot.) switch, and asymptotic performance were modeled using logistic regression, and z-scores are reported. Log-transformed response times on correct trials and total points won were modeled using linear regression, and t-values are reported. * *p < .*05; ** *p < .*01, *** *p < .*001. All models showed significant quadratic effects of age, supporting an inverse-U shaped developmental trajectory of performance.

| Performance measure (Figure) | Predictor | $\beta$ | z / t | p | sig. |
| --- | --- | --- | --- | --- | --- |
| Overall accuracy (2A) | Age (z, lin.) | 0.043 | 2.38 | 0.017 | ** |
|  | Age (z, qua.) | -0.052 | -3.11 | 0.0019 | ** |
|  | Sex | 0.009 | 0.2 | 0.77 |  |
| Total points (7A) | Age (z, lin.) | 0.003 | 0.01 | 0.99 |  |
|  | Age (z, qua.) | -1.36 | -3.11 | 0.002 | ** |
|  | Sex | 0.19 | 0.23 | 0.82 |  |
| Response times (2B) | Age (z, lin.) | -0.21 | -10.1 | < 0.001 | *** |
|  | Age (z, qua.) | 0.14 | 7.3 | < 0.001 | *** |
|  | Sex | 0.19 | 5.0 | < 0.001 | *** |
| Stay after (pot.) switch (2C) | Age (z, lin.) | 0.44 | 3.78 | < 0.001 | *** |
|  | Age (z, qua.) | -0.38 | -3.48 | < 0.001 | *** |
|  | Sex | 0.26 | 1.24 | 0.21 |  |
| Asymptotic performance (2D) | Age (z, lin.) | 0.17 | 3.57 | < 0.001 | *** |
|  | Age (z, qua.) | -0.18 | -3.97 | < 0.001 | *** |
|  | Sex | 0.030 | 0.35 | 0.73 |  |

To qualitatively assess the potential peak, without restricting the developmental trajectory to a quadratic curve, we calculated rolling performance averages (for details, see section 4.4). Most performance measures revealed peaks at around 13-15 years, including overall accuracy (Fig. 2A), points won (suppl. Fig. 7A, E), and performance after switch trials (Fig. 2C) and during stable task periods (Fig. 2D). Overall accuracy inclined steeply between ages 8-14, after which it gradually declined, settling into a stable plateau around age 20 (Fig. 2A). The willingness to repeat previous actions after a single negative outcome (Fig. 2C) showed a similarly striking increase between children and adolescents, and a (less pronounced) decline for adults. This shows that in our task, adolescents were most persistent in the face of negative feedback. Performance during stable task periods (accuracy on asymptotic trials) also was highest in adolescents, especially compared to younger participants (Fig. 2D). Response times were the only performance measure in which adolescents were outperformed by adult participants (Fig. 2B, 3D).

**Figure 2:**
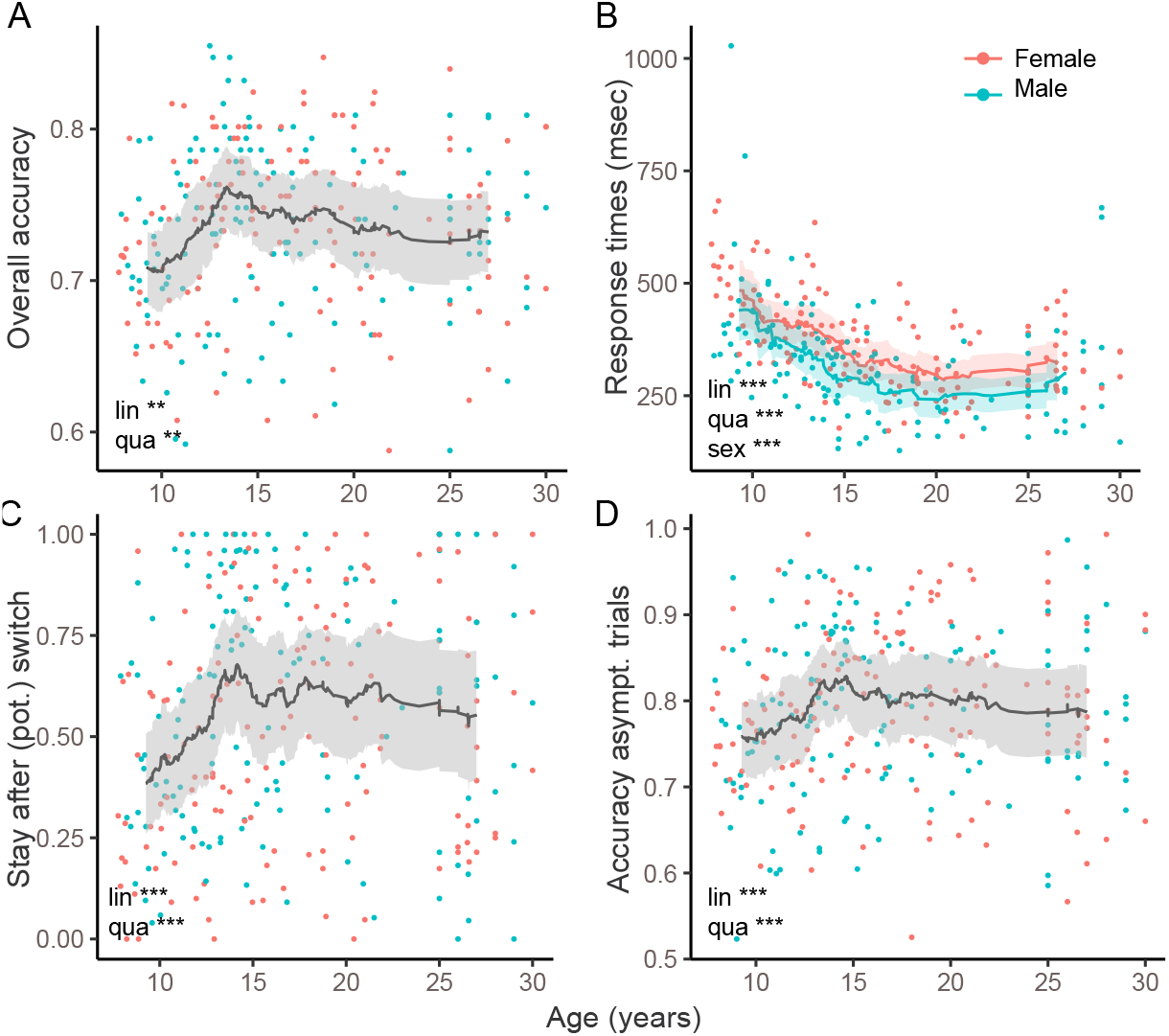
Task performance across age. Each dot shows one participant, color denotes sex. Lines show rolling averages, shades the standard error of the mean. The stars for “lin”, “qua”, and “sex” denote the significance of the effects of age, squared age, and sex on each performance measure, based on the regression models in Table 1 (* *p < .*05, ** *p < .*01, *** *p < .*001) (A) Percentage of correct choices across the entire task (120 trials), showing a peak in adolescents. The non-linear shape confirmed the significant quadratic effect of age (“qua”) on overall accuracy. (B) Median response times on correct trials. Regression coefficients differed significantly between males and females, and rolling averages are shown separately. Despite a significant quadratic effect of age, the peak for this performance measure occurred after adolescence. (C) Fraction of stay trials after (potential, “pot.”) switches (red arrows in Fig. 1C), showing an inverse U-shaped age trajectory and peak in adolescents. (D) Accuracy on asymptotic trials (grey bars in Fig. 1C), also showing an inverse U-shaped age trajectory and peak in adolescents.

**Figure 3:**
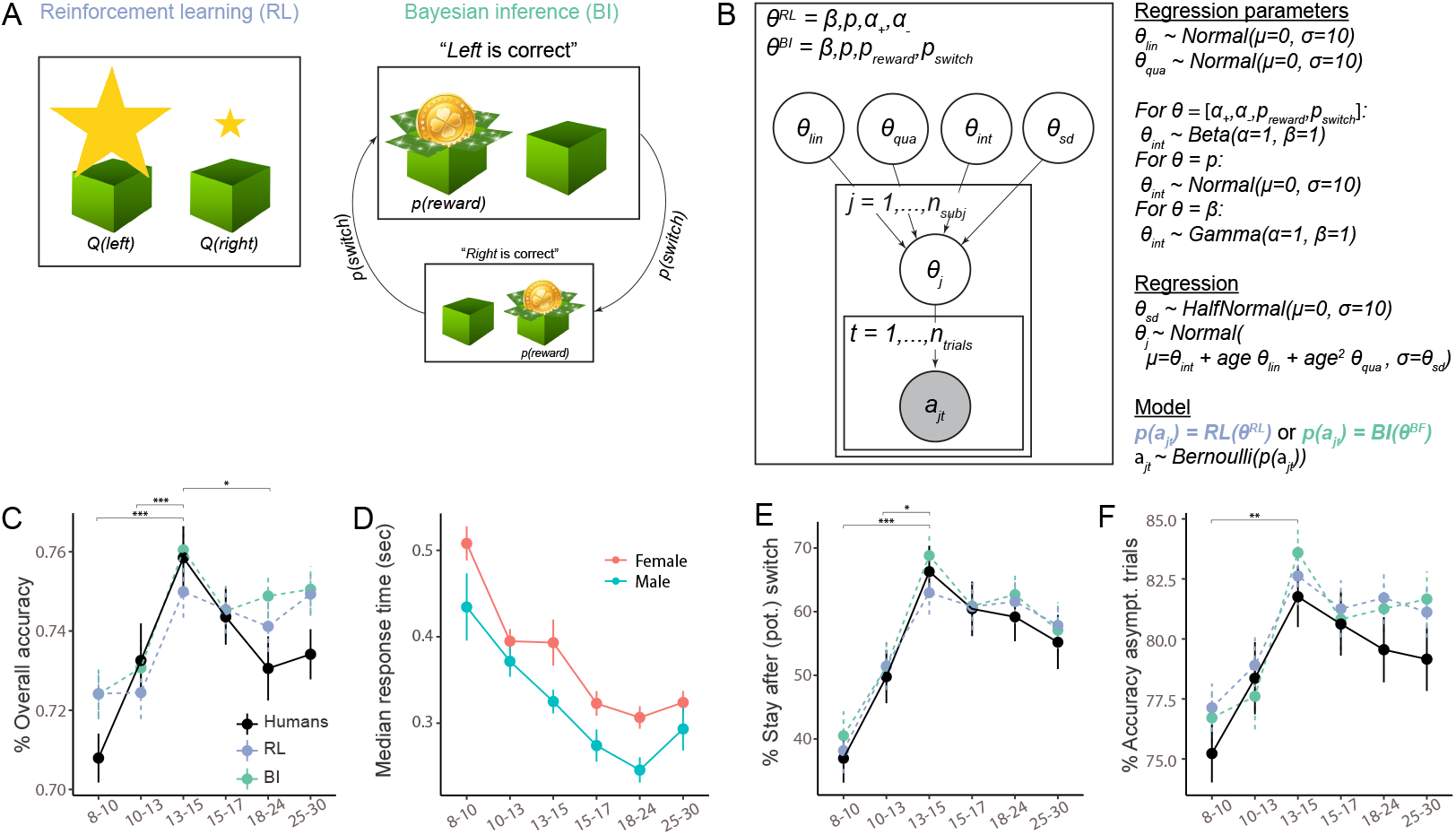
(A) Conceptual depiction of the RL and BI models. In RL (left), actions are selected based on learned values, illustrated by the size of stars (*Q*(*left*), *Q*(*right*)). In BI (right), actions are selected based on a mental model of the task, which differentiates different hidden states (“Left is correct”, “Right is correct”), and specifies the transition probability between them (*p*(*switch*)) as well as the task’s reward stochasticity (*p*(*reward*)). The sizes of the two boxes illustrate the inferred probability of being in each state. (B) Hierarchical Bayesian model fitting. Left box: RL and BI models had free parameters *θ^RL^* and *θ^BI^*, respectively. Individual parameters *θ_j_* were based on group-level parameters *θ_sd_*, *θ_int_*, *θ_lin_*, and *θ_qua_* in a regression setting (see text on the right). For each model, all parameters were simultaneously fit to the observed (shaded) sequence of actions *a_jt_* of all participants *j* and trials *t*, using MCMC sampling. Right: We chose uninformative priors for group-level parameters; the shape of each prior was based on the parameter’s allowed range. For each participant *j*, each parameter *θ* was sampled according to a linear regression model, based on group-wide standard deviation *θ_sd_*, intercept *θ_int_*, linear change with age *θ_lin_*, and quadratic change with age *θ_qua_*. Each model (RL or BI) provided a choice likelihood *p*(*a_jt_*) for each participant *j* on each trial *t*, based on individual parameters *θ_j_*. Action selection followed a Bernoulli distribution (see 4.5.3 for details). (C)-(F) Human behavior for the measures shown in Fig. 2, binned in age quantiles. (C), (E), and (F) also show simulated model behavior for model validation, verifying that models closely reproduced human behavior and age differences.

For easier visualization, we binned participants into discrete age groups, forming four equal-sized bins for participants aged 8-17, and two for adults (see section 6.2; suppl. Fig. 8A). In accordance with our hypothesis, the performance peak occurred in the intermediate age range (third youth quartile), such that adolescents between 13-15 years outperformed younger participants, older teenagers, and adults (Fig. 3C-F). Repeated, post-hoc, 5-wise Bonferroni-corrected t-tests revealed several significant differences comparing 13-to-15-year-olds to younger and older participants (Fig. 3C-F, suppl. Table 8).

We next focused on the differential effects of positive compared to negative outcomes on behavior, finding that adolescents adapted their choices more optimally to previous outcomes than younger or older participants. To show this, we used mixed-effects logistic regression to predict actions on trial *t* from predictors that encoded positive or negative outcomes on trials *t − i*, for delays 1 *≤ i ≤* 8 (for details, see section 4.4). First, we observed that the effects of positive outcomes were several times larger than the effects of negative outcomes (suppl. Table 7; Fig. 7B-F). This patterns was expected given that positive outcomes were diagnostic, whereas negative outcomes were ambivalent.

The regression model also showed an interaction between age and previous outcomes, revealing that the effects of previous outcomes on future behavior changed with age (suppl. Fig. 7B, C, E, and F; suppl. Table 7). On trials *t −* 1 and *t −* 2, positive outcomes interacted with age and squared age (all *p′s <* 0.014; suppl. Table 7), confirming that the effect of positive outcomes increased with age and then slowly plateaued (suppl. Fig. 7C, F). For negative outcomes, the signs of the interaction was opposite for trials *t −* 1 versus *t −* 2 (all *p′s <* 0.046; suppl. Table 7), showing that the effect of negative outcomes flipped, being weakest in adolescents for trial *t −* 1 (Fig. 7F), but strongest for trial *t −* 2. In other words, adolescents were best at ignoring single, ambivalent negative outcomes (*t −* 1), but most likely to integrate long-range negative outcomes (*t −* 2), which potentially indicate task switches.

To summarize, adolescents of about 13-15 years outperformed younger participants, older adolescents, and adults on a stochastic reversal task. Performance advantages were evident in several measure of task performance, and likely related to how participants responded to positive and negative outcomes. To understand better which cognitive processes underlie these patterns, we employed computational models featuring RL and BI.

### 2.3. Cognitive Modeling

We first identified a winning model of each family (RL, BI), comparing numerical fits (WAIC; Watanabe, 2013) between the most basic implementation to versions with added augmentations (suppl. Fig. 17 and Fig. 16; Table 2).

**Table 2:**
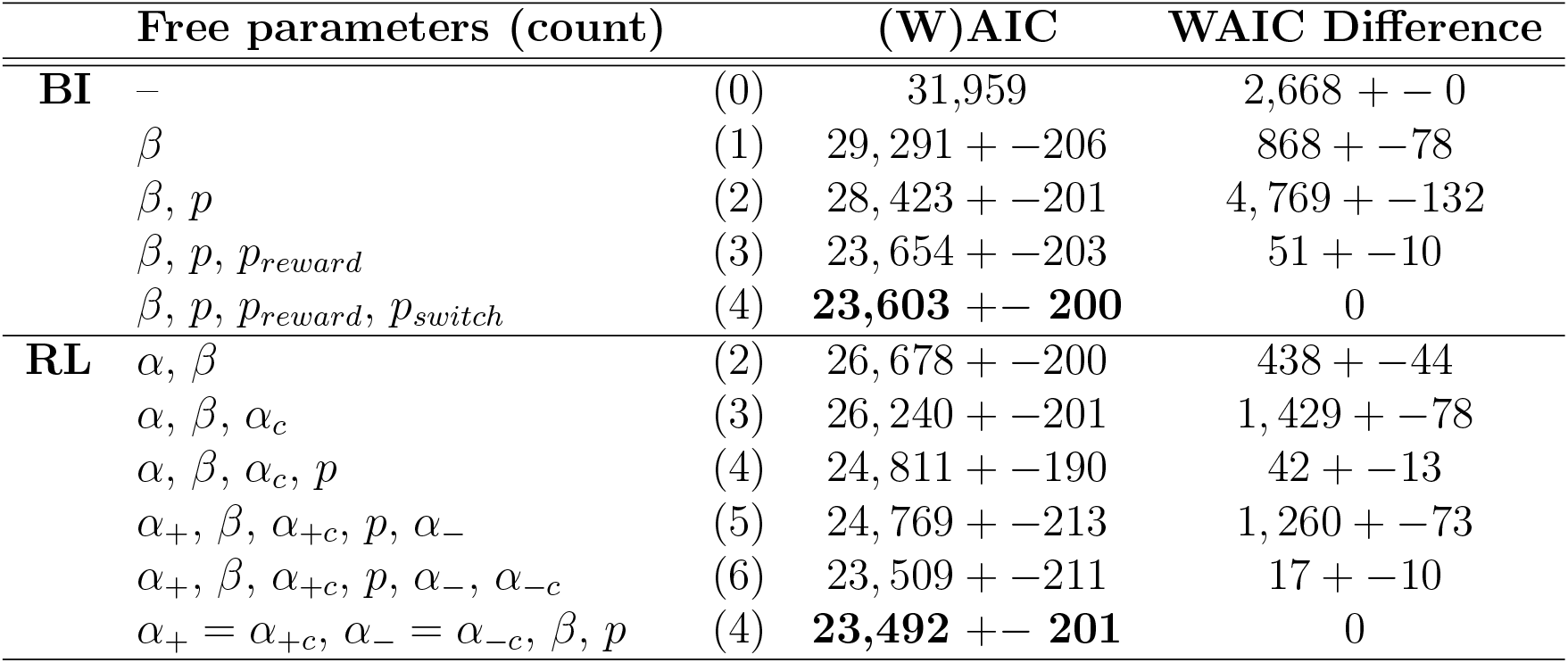
WAIC model fits and standard errors for all models, based on hierarchical Bayesian fitting. Bold numbers highlight the winning model of each class. For the parameter-free BI model, the Akaike Information Criterion (AIC) was calculated precisely. WAIC differences are relative to next-best model of the same class, and include estimated standard errors of the difference as an indicator of meaningful difference. In the RL model, “*α*” refers to the classic RL formulation in which *α*_+_ = *α_−_*. “*α_c_*” refers to the counterfactual learning rate that guides updates of unchosen actions, with *α*_+_*_c_* = *α_−c_* (see section 4.5.1).

The winning RL model had four free parameters: persistence *p*, inverse decision temperature *β*, and learning rates *α*_+_ and *α_−_* for positive and negative outcomes, respectively (section 4.5.1). In addition to “factual” action value updates on chosen actions, this model also performed “counterfactual” updates on the values of unchosen actions (Palminteri et al., 2016). For example, after receiving a reward for choosing left (factual outcome), the algorithm both decreases the value of the right choice (counterfactual update), and increases the value of the left choice (factual update). The size of counterfactual updates was controlled by learning rates *α*+ and *α_−_*, simplifying the model (Table 2). Parameters *p* and *β* controlled the translation of RL values into choices: Increasing persistence *p* increased the probability of repeating actions independently of action values. Small *β* induced decision noise (increasing exploratory choices), and large *β* allowed for reward-maximizing choices.

The winning BI model also had four parameters: besides choice-parameters *p* and *β* as in the RL model, these were task volatility *p_switch_* and reward stochasticity *p_reward_*, which characterized participants’ internal task model (Fig. 3A; section 4.5.2). *p_switch_* could represent a stable (*p_switch_* = 0) or volatile task (*p_switch_ >* 0), and *p_reward_* deterministic (*p_reward_* = 1) or stochastic outcomes (*p_reward_ <* 1). Because the actual task was based on parameters *p_switch_* = 0.05 and *p_reward_* = 0.75, an optimal agent would use these values, obtaining the most accurate inferences.

In addition to providing better model fit (Table 2), the two winning models also validated better behaviorally compared to simpler versions, closely reproducing human behavior (Palminteri et al., 2017; Wilson and Collins, 2019; Fig. 3C, E, F; suppl. Fig. 16 and Fig. 17). The winning RL model had the overall lowest WAIC score, revealing best quantitative fit, but both models validated equally well qualitatively: Both showed human-like behavior and reproduced all age differences, including adolescents’ peak in overall accuracy (Fig. 3C), proportion of staying after (potential) switch trials (Fig. 3E), asymptotic performance on non-switch trials (Fig. 3F), and their most efficient use of previous outcomes to adjust future actions (suppl. Fig. 7 D-F). Other models did not capture all these qualitative patterns (suppl. Fig. 16, Fig. 17). The closeness in WAIC scores (Table 2) and the equal ability to reproduce details of human behavior reveal that both models captured human behavior adequately, and suggest that both provide plausible explanations of the underlying cognitive processes. We therefore fitted both to participant data to estimate individual parameter values, using hierarchical Bayesian fitting (Fig. 3B; section 4.5.3).

#### 2.3.1. Age Differences in Model Parameters

Across models, three parameters showed non-monotonic age trajectories, mirroring behavioral differences: *α_−_*, *p_reward_*, and *p_switch_* declined drastically within the first three age bins (8-13 years), then reversed their trajectory and increased again, reaching slightly lower plateaus around 15 years that lasted through adulthood (Fig. 4C, G-H). For *p_switch_*, age differences were captured in a significant quadratic effect of age in the age-based model (suppl. Table 13; for detailed explanation, see section 4.5.3). For *α_−_* and *p_reward_*, differences were captured in significant pairwise differences between 13-to-15-year-olds and other age groups, tested within the age-less model (suppl. Table 12).

**Figure 4:**
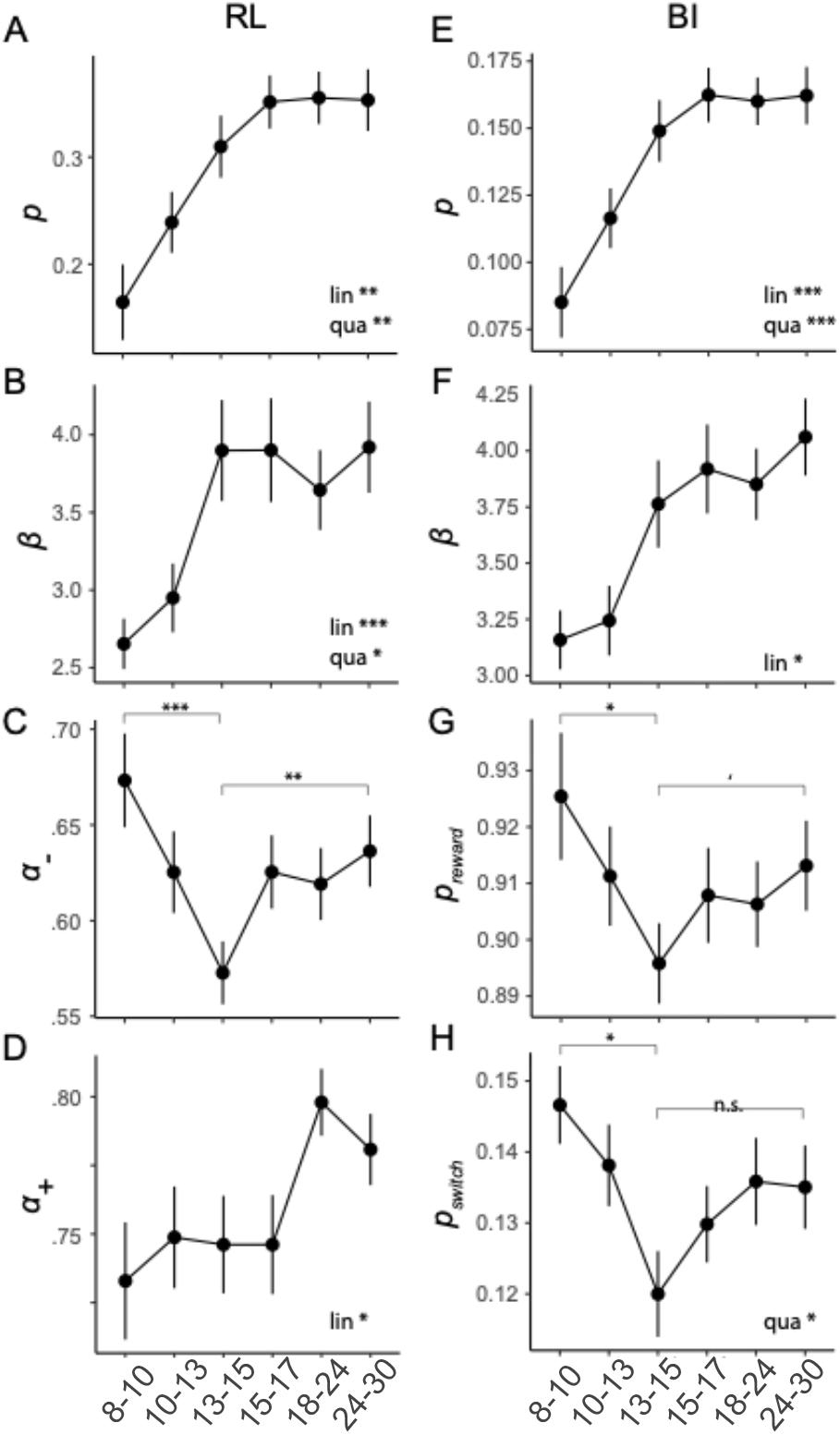
Fitted model parameters for the winning RL (left column) and BI model (right), plotted over age. Stars in combination with “lin” or “qua” indicate significant linear (“lin”) and quadratic (“qua”) effects of age on model parameters, based on the age-based fitting model. Stars on top of brackets show differences between groups, as revealed by t-tests conducted within the age-less fitting model (section 4.5.3; suppl. Tables 13 and 12). Dots (means) and error bars (standard errors) show the results of the age-less fitting model, providing an unbiased representation of individual fits. (A)-(D) RL model parameters. (E)-(H) BI model parameters.

BI’s mental model parameters *p_switch_* and *p_reward_* reflect task volatility and stochasticity (Fig. 1A), and can be compared to the true task parameters (*p_reward_* = 0.75; *p_switch_* = 0.05) to assess how optimal participants’ inferred models were. Both parameters were most optimal in 13-to-15-year-olds, whereas younger and older participants strikingly overestimated volatility (larger *p_switch_*), while underestimating stochasticity (larger *p_reward_*). Similarly in RL, *α_−_* was lowest in 13-to-15-year-olds. Indeed, lower learning rates for negative feedback *α_−_* were beneficial because they avoided premature switching based on single negative outcomes, while allowing adaptive switching after multiple negative outcomes.

In both RL and BI, choice parameters *p* and *β* increased monotonically with age, growing rapidly at first and plateauing around early adulthood (Fig. 4A, B, E, F). The age-based model (section 4.5.3) revealed that both the linear and negative quadratic effects of age were significant (suppl. Table 13). This shows that participants’ willingness to repeat previous actions independently of outcomes (*p*) and to exploit the best known option (*β*) steadily increased until adulthood, including steady growth during the teen years. Parameter *α*_+_ showed a unique stepped age trajectory, featuring relatively stable values throughout childhood and adolescence, and an increase in adults (Fig. 4D).

Through the lens of RL, these findings suggest that adolescents outperformed other age groups because they integrated negative feedback more optimally (*α_−_*). Through the lens of BI, the performance peak occurred because adolescents used a more accurate mental task model (*p_switch_* and *p_reward_*). Taken together, both models agree that behavioral differences arose from cognitive difference in the “update step” of feedback processing (i.e., value updating in RL; state inference in BI). Age differences in the “choice step” (i.e., selecting actions), however, showed monotonous age differences with steady growth during adolescents, therefore likely contributing less to the peak.

### 2.4. Integrating RL and BI—Going Beyond Specific Models

These results raise an important question: Given that both RL and BI fit human behavior well, how do we reconcile differences in their computational mechanisms? To address this, we first determined whether both models covertly employed similar computational processes, predicting the same behavior despite differences in form. A generate-and-recover analysis, however, confirmed that they truly employed different processes (Heathcote et al., 2015; Wilson and Collins, 2019; Appendix 6.3.5).

We next asked whether both models captured similar aspects of cognition by assessing how correlated parameters were between models. Parameters *p* and *β* were almost perfectly correlated between models (both *ρ >* 0.94, *p <* 0.05), suggesting high consistency between models when estimating choice processes (Fig. 5B). Parameter *p_reward_* (BI) was strongly correlated with *α_−_* (RL), suggesting that beliefs about task stochasticity and learning rates for negative outcomes played similar roles across models, presumably in participants’ response to negative outcomes. The other mental-model parameter, *p_switch_* (BI), was strongly negatively correlated with *β* (RL), suggesting that beliefs about task volatility in the BI model captured aspects that were explained by decision noise in the RL model. This is consistent with the observation that an agent that expects high volatility could be mistaken for one that acts with large noise, given that both will make choices that are inconsistent with previous outcomes. The only parameter that showed no large correlations with other parameters was *α*_+_ (RL), potentially reflecting a cognitive process uniquely captured by RL. Taken together, some parameters likely captured similar cognitive processes in both models, despite differences in their functional form, shown by large correlations between models. Other parameters were more unique, potentially reflecting model-specific cognitive processes. Further analyses confirmed high shared explained variance between both models, using multiple regression (section 6.3.7).

**Figure 5:**
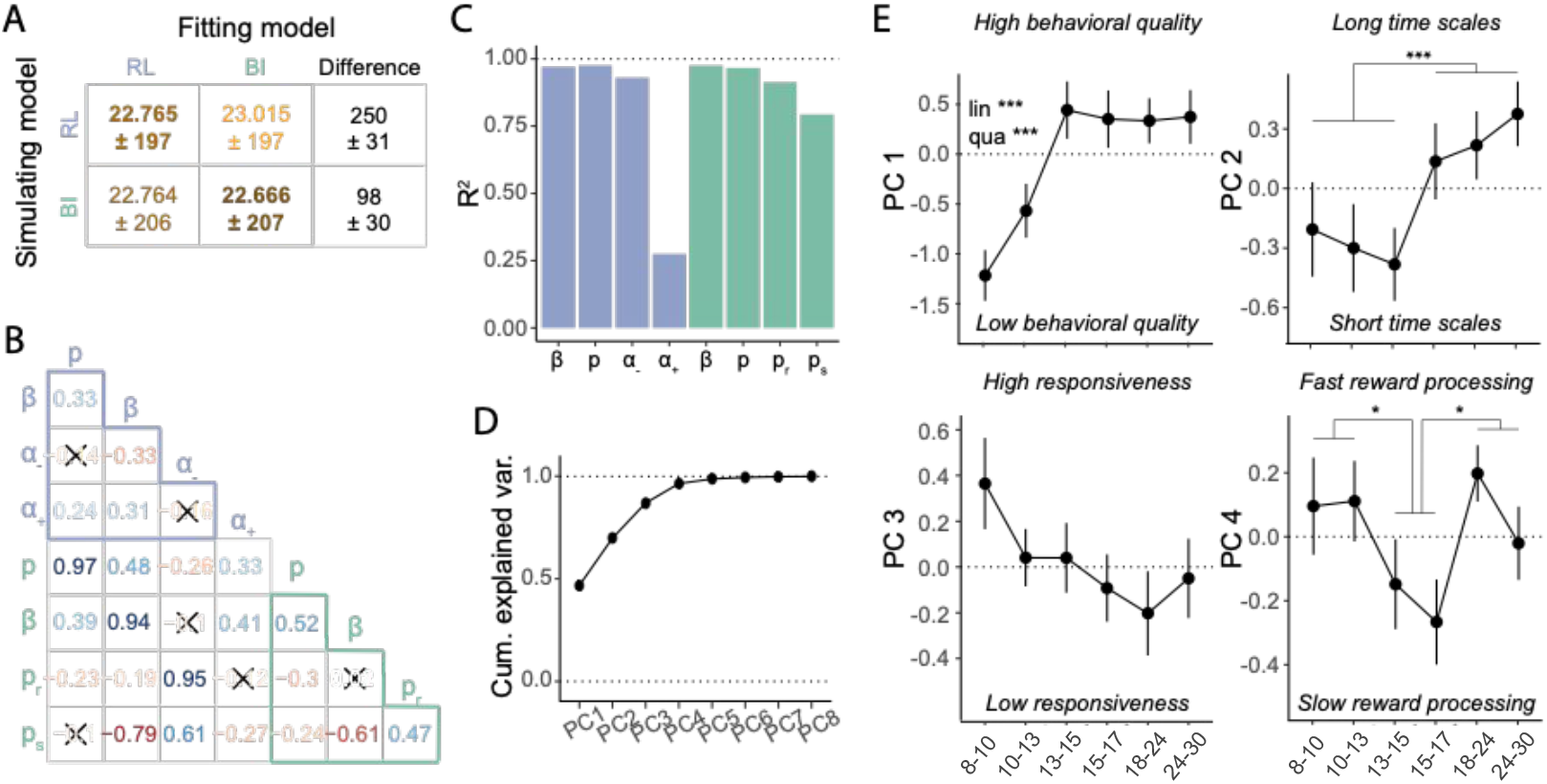
Relating RL and BI models. (A) Model recovery. WAIC scores were worse (larger; lighter colors) when recovering behavior that was simulated from one model (row) using the other model (column), than when using the same model (diagonal), revealing that the models were discriminable. The difference in fit was smaller for BI simulations (bottom row), suggesting that the RL model captured BI behavior better than the other way around (top row). (B) Spearman pairwise correlations between model parameters. Red (blue) hue indicates negative (positive) correlation, saturation indicates correlation strength. Non-significant correlations are crossed out (Bonferroni-corrected at *p* = 0.00089). Light-blue (teal) letters refer to RL (BI) model parameters. Light-blue / teal-colored triangles show correlations within each model, remaining cells show correlations between models. (C) Variance of each parameter explained by parameters and interactions of the other model (“*R*^2^”), estimated through linear regression. All four BI parameters (green) were predicted almost perfectly by the RL parameters, and all RL parameters except for *α*_+_ (RL) were predicted by the BI parameters. (D)-(E) Results of PCA on model parameters. (D) Cumulative variance explained by all principal components PC1-8. The first four components captured 96.5% of total parameter variance. (E) Age-related differences in PC1-4: PC1 reflected overall behavioral quality and showed rapid development between ages 8-13, which were captured by linear (“lin”) and quadratic (“qua”) effects in a regression model. PC2 captured a step-like transition from shorter to longer updating time scales at age 15, as revealed by PC-based model simulations (Supplements). PC3 showed no significant age effects. PC4 captured the variance in *α*_+_ and differed between adolescents 15-17 and both 8-13 year olds and adults. PC2 and PC4 were analyzed using t-tests. * *p < .*05; ** *p < .*01, *** *p < .*001.

So far, we have provided two separate cognitive explanations for why adolescents performed better than other age groups: RL poses differences in value learning as the main driver, whereas BI poses differences in mental model-based inference. Could a single, broader explanation combine these insights and provide more general understanding of adolescent cognitive processing? To test this, we used PCA to unveil the lower-dimensional structure embedded in the 8-dimensional parameter space created by both models (for details, see section 4.5.5). We found that the PCA’s first four principle components (PCs) explained almost all variance (96.5%; Fig. 5D), showing that individual differences in all 8 model parameters could be summarized by just 4 abstract cognitive dimensions, which distill the insights of both models while abstracting away redundancies. To understand what these abstract dimensions reflected, we used a simulation-based approach that took advantage of the fact that each PC was a linear combination of the original model parameters (Table 14), such that we could directly simulate effects of PCs on behavior using our computational models.

This analysis revealed that PC1, capturing the largest proportion of parameter variance, reflected a broad measure of behavioral quality; PC2 represented integration time scales; PC3 captured responsiveness to task outcomes; and PC4 uniquely captured RL parameter *α*_+_. A detailed description of each PC is provided in supplement 6.3.8. Three of these four PCs (PC1, PC2, and PC4) showed prominent age effects: PC1 (behavioral quality) increased drastically until age 13, at which it reached a stable plateau that lasted—unchanged—throughout adulthood (Fig. 5E, top-left). Regression models revealed significant linear and quadratic effects of age on PC1 (lin.: *β* = *−*0.47, *t* = *−*4.0, *p <* 0.001; quad.: *β* = 0.011, *t* = 3.43, *p <* 0.001), with no effect of sex (*β* = 0.020, *t* = 0.091, *p* = 0.93). This suggests that the left side of the U-shaped trajectory in task performance (Fig. 2; suppl. Fig. 7; Fig. 3C-F) might be caused by the development of behavioral quality (PC1): The peak in 13-to-15-year-olds compared to younger participants could be explained by the fact that 13-to-15-year-olds had already reached adult levels of behavioral quality, while younger participants showed noisier, less focused, and less consistent behavior.

By contrast, PC2 (updating time scales) followed a step function, such that participants in the three youngest age bins (8-15 years) acted on shorter times scales than participants in the three oldest bins (15-30; Fig. 5E, top-right; post-hoc t-test comparing both groups: *t*(266.2) = 3.44, *p <* 0.001). This pattern is in accordance with the interpretation that children’s shorter time scales, facilitating rapid behavioral switches (suppl. Fig. 19B, left), were more beneficial for the current task than adults’ longer time scales, which impeded switching (suppl. Fig. 19B, right). Differences in subjective time scale might therefore be the determining factor that allowed adolescents to outperform older participants, including adults.

PC4 (positive updates) differentiated the two adolescent age bins (13-17) from both younger (8-13) and older (18-30) participants (Fig. 5E, bottom-right), as revealed by significant post-hoc, Bonferroni-corrected, t-tests (8-13 vs 13-17: *t*(176.8) = 2.28, *p* = 0.047; 13-17 vs 18-30: *t*(176.6) = 2.49, *p* = 0.028). In other words, after accounting for variance in PC1-PC3, the remaining variance was explained by 13-to-17-year-olds’ relatively longer updating timescales for positive outcomes (positive outcomes had relatively weaker immediate, but stronger long-lasting effects). In sum, the PCA revealed four dimensions that combine the findings of both computational models, potentially allowing for model-independent insights into developmental cognitive differences: Adolescents’ unique competence in our task might be result of adult-like behavioral quality in combination with child-like time scales and unique adolescent processing of positive feedback.

## 3. Discussion

Across species, the adolescent transition brings great challenges for learning and exploration, which may have caused the adolescent brain to evolve behavioral tendencies that promote adaptive learning in rapidly changing, uncertain environments (Dahl et al., 2018). To test this idea, we examined the choice behavior of a large sample across a wide age range in a volatile and stochastic reversal task adapted from rodent studies (Tai et al., 2012). This research fills a knowledge gap regarding the adolescent development of reversal learning (also see Hauser et al., 2015; Javadi et al., 2014; van der Schaaf et al., 2011), inspired by rapidly-expanding research highlighting the developmentally-unique role of adolescence across socio-emotional and cognitive contexts (Dahl et al., 2018; DePasque and Galván, 2017; Lloyd et al., 2020; Lourenco and Casey, 2013; Sercombe, 2014), and by the non-linear development of neural and endocrine systems underlying reversal learning (Blakemore et al., 2010; Braams et al., 2015; Giedd et al., 1999; Piekarski, Johnson, et al., 2017; Somerville and Casey, 2010; Sowell et al., 2003; Toga et al., 2006).

### 3.1. Summary of Findings

We observed an adolescent peak in performance, which was evident in adolescents’ highest overall accuracy (Fig. 2A) and winning most points (suppl. Fig. 7A, E). This peak was associated with adolescents’ increased willingness to ignore non-diagnostic negative feedback (Fig. 2C) and to show persistent choices during stable task periods (Fig. 2D). Adolescents used negative feedback most optimally to guide future choices, being least affected by proximal, but most sensitive to distal outcomes (suppl. Fig. 7C, D, G, H). These findings support our prediction that adolescents make better decisions in volatile and stochastic environments, potentially due to differences in negative feedback processing, in accordance with prior research that has shown unique feedback processing in adolescents (e.g., Christakou et al., 2013; Davidow et al., 2016; Palminteri et al., 2016; van den Bos et al., 2009; for review, see Lourenco and Casey, 2013).

Which cognitive processes underlie this performance advantage? Adolescents might learn at different speeds than younger or older participants, as suggested, e.g., by Davidow et al., 2016, or might process particular feedback types differently (e.g., Palminteri et al., 2016). These hypotheses can be tested using computational modeling in the RL framework, which explicitly estimates learning rates for different kinds of feedback.

It is also possible, however, that adolescents outperformed other participants due to a better understanding of the task dynamics, which would allow them to predict more accurately whether a switch had occurred, for example. Indeed, others have argued that both “model-based” behavior (Decker et al., 2016) and the tendency to employ counterfactual reasoning (Palminteri et al., 2016) increase with age, in accordance with age differences in mental task models. This hypothesis can be tested using computational modeling in the BI framework, which explicitly estimates the parameters of participants’ mental models and inference processes.

Furthermore, adolescents might explore differently (Gopnik et al., 2017; Lloyd et al., 2020; Somerville et al., 2017) or might be more persistent, a behavioral pattern commonly linked to the PFC (Kehagia et al., 2010; Morris et al., 2016), which continues maturation during adolescence (DePasque and Galván, 2017; Giedd et al., 1999; Toga et al., 2006). Whereas the previous hypotheses targeted the “updating” step of decision making, these two concern the “choice” step, and can be tested in both RL and BI frameworks. Our study revealed that several explanations exist for adolescents’ superior performance: The RL model showed reduced learning speeds for negative outcomes (Fig. 4C), supporting the hypothesis in terms of differential feedback responses. The BI model suggested improved mental models, supporting the hypothesis about differences in mental models and inference (Fig. 4G, H). Crucially, the quantitative fit of both models to human data was similar (Table 2), and they both qualitatively reproduced human behavior in simulation (Fig. 3), suggesting that both explanations are valid. Furthermore, both models agreed on developmental differences in exploration/exploitation and persistence, as suggested by the last hypotheses. However, these differences were unlikely the cause for the adolescent advantage because they showed monotonic trajectories between childhood and adulthood (Fig. 4A, B, E, G), rather than an adolescent peak. Taken together, our study suggests that adolescents make better decisions in stochastic and volatile environments than younger or older people, due to non-monotonic age differences in negative feedback processing and mental model accuracy, which peak during adolescence.

Both explanations, however, are framed within a specific computational model. Can we draw more general conclusions? Combining the unique insights of each model while stripping away redundancies, our PCA investigation revealed that developmental changes might be captured by three abstract, model-independent dimensions that vary with age: behavioral quality (PC1), time scales (PC2), and reward processing (PC4). Behavioral quality—likely capturing sufficient understanding of the task and experimental context, participant compliance, attentional focus, etc.—reached adult levels in early adolescence and showed no more age-related differences thereafter. Time scales, on the other hand—likely capturing an extended planning horizon, long-term credit assignment, memory, prolonged attention, etc.— only started to increase during late adolescence, in accordance with our behavioral measure of flexibility (6.3.1). Finally, reward processing was slower during adolescence compared to younger or older ages. Taken together, adolescents’ behavioral advantage might be a combination of already adult-like quality of behavior, still child-like time scales, and unique reward processing.

### 3.2. Setting or Adaptation?

These findings can be interpreted in two ways (Nussenbaum and Hartley, 2019): 1) Based on a *settings* account, adolescents integrate negative feedback more slowly than other age groups (*α_−_*), expect fewer rewards (*p_reward_*) and less volatility (*p_switch_*), and achieve adult-like behavioral quality (PC1), but child-like short time scales (PC2) and slow reward processing (PC4). These “settings” are developmentally fixed, i.e., expected to guide behavior across experiments and real-life situations. 2) The *adaptation* account, on the other hand, states that adolescents chose the most appropriate cognitive settings specifically for the current task, and might have chosen different settings in different contexts. Our results therefore highlight adolescents’ adaptability to volatile and stochastic environments.

A recent review (Nussenbaum and Hartley, 2019) showed favorable empirical evidence for the adaptation account compared to settings, given that specific parameter values differ widely between studies, whereas parameter adaptiveness is more consistent (also see Eckstein, Master, et al., 2021; Eckstein, Wilbrecht, et al., 2021). Another argument for adaption is that adolescents exhibited *balanced* learning in a previous study (van der Schaaf et al., 2011), responding similarly to rewards and punishment (Fig. 3A; children and adults responded more strongly to punishment and rewards, respectively). In our study, however, adolescents exhibited the most *imbalanced* learning of all age groups, responding least strongly to negative feedback. This shows a contradiction between both studies based on a settings view. However, both studies agree in that adolescents adapted best to the specific task demands, supporting an adaptation-based view: In van der Schaaf et al., 2011, both positive and negative outcomes were diagnostic, requiring balanced learning, whereas in our study, only positive outcomes were diagnostic, requiring imbalanced learning.

Taken together, the specific parameter values obtained in this study likely shed less light on specific adolescent behavioral tendencies related to negative feedback processing, prior expectations about environmental volatility and stochasticity, etc., but showcase the increased ability to quickly and effortlessly adapt to stochastic and volatile tasks.

### 3.3. General Cognitive Abilities

A caveat of our study is the use of a cross-sectional rather than longitudinal design. We cannot exclude, for example, that adolescents had higher IQ scores, better schooling, or higher socio-economic status than participants of other ages. If this was the case, the performance peak in adolescence might reflect a difference in task-unrelated factors rather than unique adaptation to stochasticity and volatility. However, several arguments speak against this possibility, including recruitment procedures, supplementary analyses, and the distinctness of the U-shaped pattern observed in this task compared to the linear trajectories observed in other tasks performed by the same sample (see section 6.4.1).

### 3.4. A Role of Puberty?

Despite showing specific age-related differences, our study does not elucidate which biological mechanisms underlie these. There is growing evidence that gonadal hormones affect inhibitory neurotransmission, spine pruning, and other variables in the prefrontal cortex of rodents (Delevich et al., 2019; Delevich et al., 2018; Drzewiecki et al., 2016; Juraska and Willing, 2017; Piekarski, Boivin, et al., 2017; Piekarski, Johnson, et al., 2017), and evidence for puberty-related neurobehavioral change is also accumulating in human studies (Blakemore et al., 2010; Braams et al., 2015; Gracia-Tabuenca et al., 2021; Laube, van den Bos, et al., 2020; Op de Macks et al., 2016), suggesting that puberty-related changes in brain chemistry might be a mechanism behind the observed differences. We assessed pubertal status and investigated its role in the developmental changes we observed (see section 6.3.3). While some trends emerged that deserve more detailed investigation in future research, particularly with regard to early puberty, our study was inconclusive on this issue.

### 3.5. Dual-Model Approach to Cognitive Modeling

Basic RL and BI (as described in section 1.3) employ different cognitive mechanisms (see sections 4.5.1 and 4.5.2) and predict different behaviors on our task (suppl. Fig. 16 and 17), justifying their combined use to gain additive insights. However, we augmented each model to approximate humans, leading to more similar behavior—and potentially overlapping cognitive mechanisms. Is this a problem for our dual-model approach?

Two arguments justify the approach: 1) Both models *explain* the cognitive process differently. Whereas RL explains it in terms of learning and differentiation of outcome types, BI explains it in terms of mental-model based predictions and inference. Hence, invoking different cognitive concepts, both explanations are non-redundant and provide additive insights. 2) Both models also *differ* in meaningful ways, both behaviorally (Fig. 5A; suppl. Fig. 18; suppl. section 6.3.6) and in terms of the cognitive processes captured by model parameters (Fig. 5B and C). This implies that both models capture different aspects of human cognitive processing, providing additive insights. Taking a step back, the most common computational modeling approach selects a family of candidate models (e.g., RL) and identifies the best-fitting one, interpreting it as the cognitive process employed by participants. An issue with this approach is that a model from a different family (e.g., BI) might provide a better fit than any of the tested models. To address this issue, we fitted models of multiple families, ensuring large coverage of the space of cognitive hypotheses.

However, a difficulty with our approach is that in addition to quantitative criteria of model fit (e.g., fit, complexity; Bayes factor, AIC; Mulder and Wagenmakers, 2016; Pitt and Myung, 2002; Watanabe, 2013), qualitative criteria become increasingly important (e.g., interpretability, appropriateness for current hypotheses, conciseness, generality; Blohm et al., 2020; Kording et al., 2020; Uttal, 1990; Webb, 2001). However, qualitative criteria are more difficult to assess because they depend on scientific goals (e.g., explanation versus prediction; Bernardo and Smith, 2009; Navarro, 2019) and research philosophy (Blohm et al., 2020). Furthermore, qualitative and quantitative criteria can be at odds, inconveniencing model selection (Jacobs and Grainger, 1994). To alleviate these issues, we focused on a range of criteria, including numerical fit (WAIC; slight advantage for RL), reproduction of participant behavior (equally good), continuity with previous neuroscientific research (RL), link to specific neural pathways (RL), centrality for developmental research (equal), claimed superiority in current paradigm (BI), and interpretability (BI: model parameters map directly onto main concepts *p_switch_*: stochasticity, *p_reward_*: volatility). Because this survey did not produce a clear winner, and both models fitted excellently without being redundant, we opted to select two winners. This provided the benefits of *converging evidence* (e.g., replication: *β_RL_ ↔ β_BI_*, *p_RL_ ↔ p_BI_*; parallelism between models: *p_reward_ ↔ α_−_*), *distinct insights* (e.g., RL: importance of learning, differential processing of feedback types; BI: importance of inference, mental models), and the possibility to *combine* both models to expose more abstract factors (PC1, PC2, PC4) that differentiate adolescent cognitive processing from younger and older participants.

### 3.6. Conclusion

In conclusion, we showed that adolescents outperformed younger participants and adults in a volatile and uncertain context, two factors that might have specific relevance in the transition of adolescence. We used two computational models to examine the cognitive processes underlying this development, RL and BI. These models suggested that adolescents achieved better performance for different reasons: (1) They were best at accurately assessing the volatility and stochasticity of the environment, and integrated negative outcomes most appropriately (U-shapes in *p_reward_*, *p_switch_*, and *α_−_*). (2) They combined adult-like behavioral quality (PC1), child-like time scales (PC2), and developmentally-unique processing of positive outcomes (PC4). Pubertal development and steroid hormones may impact a subset of these processes, yet causality is difficult to determine without manipulation or longitudinal designs (Kraemer et al., 2000).

For purposes of translation from the lab to the “real world”, our study indicates that how youth learn and decide changes in a nonlinear fashion as they grow. This underscores the importance of youth-serving programs that are developmentally informed and avoid a one-size-fits-all approach. Finally, these data support a positive view of adolescence and the idea that the adolescent brain exhibits remarkable learning capacities that should be celebrated.

## 4. Methods

### 4.1. Participants

All procedures were approved by the Committee for the Protection of Human Subjects at the University of California, Berkeley. We tested 312 participants: 191 children and adolescents (ages 8-17) and 55 adults (ages 25-30) were recruited from the community, using online ads (e.g., on neighborhood forums), flyers at community events (e.g., local farmers markets), and physicals posts in the neighborhood (e.g., printed ads). Community participants completed a battery of computerized tasks, questionnaires, and saliva samples (Master et al., 2020). In addition, 66 university undergraduate students (aged 18-50) were recruited through UC Berkeley’s Research Participation Pool, and completed the same four tasks, but not the pubertal-development questionnaire (PDS; Petersen et al., 1988) or saliva sample. Community participants were prescreened for the absence of present or past psychological and neurological disorders; the undergraduate sample indicated the absence of these. Community participants were compensated with 25$ for the 1-2 hour in-lab portion of the experiment and 25$ for completing optional take-home saliva samples; undergraduate students received course credit for participation.

#### Exclusion Criteria

Out of the 191 participants under 18, 184 completed the current task; reasons for not completing the task included getting tired, running out of time, and technical issues. Five participants (mean age 10.0 years) were excluded because their mean accuracy was below 58% (chance: 50%), an elbow point in accuracy, which suggests they did not pay attention to the task. This led to a sample of 179 participants under 18 (male: 96, female: 83). Two participants from the undergraduate sample were excluded because they were older than 30, leading to a sample aged 18-28; 7 were excluded because they failed to indicate their age. This led to a final sample of 57 undergraduate participants (male: 19, female: 38). All 55 adult community participants (male: 26, female: 29) completed the task and were included in the analyses, leading to a sample size of 179 participants below 18, and 291 in total (suppl. Fig. 8).

### 4.2. Testing Procedure

After entering the testing room, participants under 18 years and their guardians provided informed assent and permission, respectively; participants over 18 provided informed consent. Guardians and participants over 18 filled out a demographic form. Participants were led into a quiet testing room in view of their guardians, where they used a video game controller to complete four computerized tasks (for more details about the other tasks, see Eckstein, Master, et al., 2021; Master et al., 2020; Xia et al., 2020; for a comparison of all tasks, see Eckstein, Master, et al., 2021; Eckstein, Wilbrecht, et al., 2021). At the conclusion of the tasks, participants between 11 and 18 completed the PDS questionnaire, were measured in height and weight, and compensated with $25 Amazon gift cards. The entire session took 2-3 hours for community participants (e.g., some younger participants took more breaks), and 1 hour for undergraduate participants (who did not complete the puberty measures and saliva sample). We paid great attention to the fact that participants took sufficient breaks between tasks to avoid excessive fatigue and limit the effects of the differences in testing duration.

### 4.3. Task Design

The goal of the task was to collect golden coins, which were hidden in one of two boxes. On each trial, participants decided which box to open, and either received a reward (coin) or not (empty). Task contingencies— i.e., which box was correct and therefore able to produce coins—switched unpredictably throughout the task (Fig. 1B). Before the main task, participants completed a 3-step tutorial: 1) A prompt explained that only one of the boxes contained a coin (was “magical”), and participants completed 10 practice trials on which one box was always rewarded and the other never (deterministic phase). 2) Another prompt explained that the magical box sometimes switches sides, and participants received 8 trials on which only second box was rewarded, followed by 8 trials on which only the first box was rewarded (switching phase). 3) The last prompt explained that the magical box did not always contain a coin, and and led into the main task with 120 trials.

In the main task, the correct box was rewarded in 75% of trials; the incorrect box was never rewarded. After participants reached a performance criterion, it became possible for contingencies to switch (without notice), such that the previously incorrect box became the correct one. The performance criterion was to collect 7-15 rewards, with the specific number pre-randomized for each block (any number of non-rewarded trials was allowed in-between rewarded trials). Switches only occurred after rewarded trials, and the first correct choice after a switch was always rewarded (while retaining an average of 75% probability of reward for correct choices), for consistency with the rodent task (Tai et al., 2012).

### 4.4. Behavioral Analyses

We calculated age-based rolling performance averages by averaging the mean performance of 50 subsequent participants ordered by age. Standard errors were calculated in the same way.

We assessed the effects of age on behavioral outcomes (Fig. 2), using (logistic) mixed-effects regression models using the package lme4 (Bates et al., 2015) in R (RCoreTeam, 2016). All models included the following regressors to predict outcomes (e.g., overall accuracy, response times): Z-scored age, to assess the linear effect of age on the outcome; squared, z-scored age, to assess the quadratic (U-shaped) effect of age; and sex; furthermore, all models specified random effects of participants, allowing participants’ intercepts and slopes to vary independently. Additional predictors are noted in the main text.

We assessed the effects of previous outcomes on participants’ choices (suppl. Fig. 7B, C, E, F) using logistic mixed-effects regression, predicting actions (left, right) from previous outcomes (details below), while testing for effects of and interactions with sex, z-scored age, and z-scored quadratic age, specifying participants as mixed effects. We included one predictor for positive and one for negative outcomes at each delay *i* with respect to the predicted action (e.g., *i* = 1 trial ago). Outcome predictors were coded −1 for left and +1 for right choices (0 otherwise). Including predictors of trials 1 *i* 8 provided the best model fit (suppl. Table 7). To visualize the results of this model including all participants, we also ran separate models for each participant (suppl. Fig. 7B, C, E, F).

### 4.5. Computational Models

#### 4.5.1. Reinforcement Learning (RL) Models

A basic RL model has two parameters, learning rate *α* and decision temperature *β*. On each trial *t*, the value *Q_t_*(*a*) of action *a* is updated based on the observed outcome *o_t_ ∈* [0, 1] (no reward, reward):

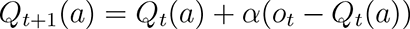

Action values inform choices probabilistically, based on a softmax transformation:

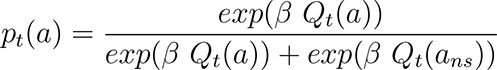

Here, *a* is the selected, and *a_ns_* the non-selected action.

Compared to this basic 2-parameter model, the best-fit 4-parameter model was augmented by splitting learning rates into *α*_+_ and *α_−_*, adding persistence parameter *p*, and the ability for counterfactual updating. We explain each in turn: Splitting learning rates allowed to differentiate updates for rewarded (*o_t_* = 1) versus non-rewarded (*o_t_* = 0) trials, with independent *α_−_* and *α*_+_:

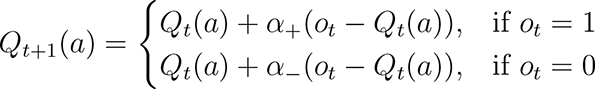

Choice persistence or “stickiness” *p* changed the value of the previously-selected action *a_t_* on the subsequent trial, biasing toward staying (*p >* 0) or switching (*p <* 0):

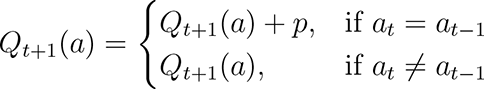

Counterfactual updating allows updates to non-selected actions based on counterfactual outcomes 1 *− o_t_*:

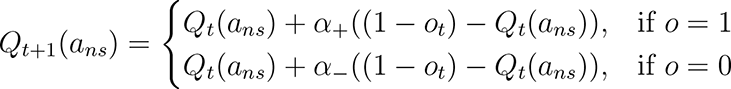

Initially, we used four parameters *α*_+_, *α*_+_*_c_*, *α_−_*, and *α_−c_* to represent each combination of value-based (“+” versus “-”) and counter-factual (“c”) versus factual updating, but collapsing *α*_+_ = *α*_+_*_c_* and *α_−_* = *α_−c_* improved model fit (Table 2). This suggests that outcomes triggered equal-sized updates to chosen and unchosen actions.

This final model can be interpreted as basing decisions on a single value estimate (value difference between both actions), rather than independent value estimates for each action because chosen and unchosen actions were updated to the same degree and in opposite directions on each trial. Action values were initialized at 0.5 for all models.

#### 4.5.2. Bayesian Inference (BI) Models

The BI model is based on two hidden states: “Left action is correct” (*a_left_* = *cor*) and “Right action is correct” (*a_right_* = *cor*). On each trial, the hidden state switches with probability *p_switch_*. In each state, the probability of receiving a reward for the correct action is *p_reward_* (Fig. 3A). On each trial, actions are selected in two phases, using a Bayesian Filter algorithm (Sarkka, 2013): (1) In the *estimation phase*, the hidden state of the previous trial *t −* 1 is inferred based on outcome *o_t−_*_1_, using Bayes rule:

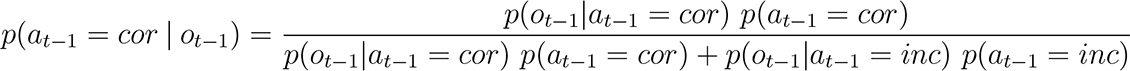

*p*(*a_t−_*_1_ = *cor*) is the prior probability that *a_t−_*_1_ is correct (on the first trial, *p*(*a* = *cor*) = 0.5 for *a_left_* and *a_right_*). *p*(*o_t−_*_1_*|a_t−_*_1_) is the likelihood of the observed outcome *o_t−_*_1_ given action *a_t−_*_1_. Likelihoods are (dropping underscripts for clarity): *p*(*o* = 1*|a* = *cor*) = *p_reward_*, *p*(*o* = 0*|a* = *cor*) = 1 *p_reward_*, *p*(*o* = 1 *a* = *inc*) = *E*, and *p*(*o* = 0 *a* = *cor*) = 1 *E*. *E* is the probability of receiving a reward for an incorrect action, which was 0 in reality, but set to *E* = 0.0001 to avoid model degeneracy.

(2) In the *prediction phase*, the possibility of state switches is taken into account by propagating the inferred hidden-state belief at *t* − 1 forward to trial *t*:

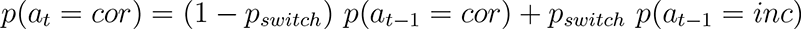

We first assessed a parameter-free version of the BI model, truthfully setting *p_reward_* = 0.75, and *p_switch_* = 0.05. Lacking free parameters, this model was unable to capture individual differences and led to poor qualitative (suppl. Fig. 17A) and quantitative model fit (Table 2). The best-fit BI model had four free parameters: *p_reward_* and *p_switch_*, as well as the choice parameters *β* and *p*, like the winning RL model. *β* and *p* were introduced by applying a softmax to *p*(*a_t_* = *cor*) to calculate *p*(*a_t_*), the probability of selecting action *a* on trial *t*:

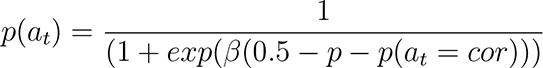

When both actions had the same probability and persistence *p >* 0, then staying was more likely; when *p <* 0, then switching was more likely.

#### 4.5.3. Model Fitting and Comparison

We fitted parameters using hierarchical Bayesian methods (Katahira, 2016; M. D. Lee, 2011; van den Bos et al., 2017; Fig. 3B), whose parameter recovery clearly superseded those of classical maximum-likelihood fitting (suppl. Fig. 6). Rather than fitting individual participants, hierarchical Bayesian model fitting estimates the parameters of a population jointly by maximizing the posterior probability *p*(*θ data*) of all parameters *θ* conditioned on the observed *data*, using Bayesian inference:

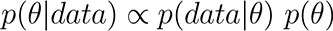

An advantage of hierarchical Bayesian model fitting is that individual parameters are embedded in a hierarchical structure of priors, which helps resolve uncertainty at the individual level.

We ran two models to fit parameters: The “age-less” model was used to estimate participants’ parameters in a non-biased way and conduct binned analyses on parameter differences; the “age-based” model was used to statistically assess the shapes of parameters’ age trajectories. In the age-less model, each individual *j*’s parameters 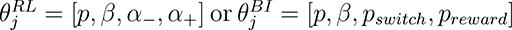 were drawn from group-based prior parameter distributions. Parameters were drawn from appropriately-shaped prior distributions, limiting ranges where necessary, which where based on non-informative, appropriate hyperpriors (suppl. Table 5).

Next, we fitted the model by determining the group-level and individual parameters with the largest posterior probability under the behavioral data *p*(*θ data*). Because *p*(*θ data*) was analytically intractable, we approximated it using Markov-Chain Monte Carlo sampling, using the no-U-Turn sampler from the PyMC3 package in python (Salvatier et al., 2016). We ran 2 chains per model with 6,000 samples per chain, discarding the first 1,000 as burnin. All models converged with small MC errors, sufficient effective sample sizes, and *R*^^^ close to 1 (suppl. Table 6). For model comparison, we used the Watanabe-Akaike information criterion (WAIC), which estimates the expected out-of-sample prediction error using a bias-corrected adjustment of within-sample error (Watanabe, 2013).

To obtain participants’ individual fitted parameters, we calculated the means over all posterior samples (Fig. 4, suppl. Figures 15, 16, and 17). To test whether a parameter *θ* differed between two age groups *a*1 and *a*2, we determined the number of MCMC samples in which the parameter was larger in one group than the other, i.e., the expectation E(*θ_a_*_1_ *< θ_a_*_2_) across MCMC samples. *p <* 0.05 was used to determine significance. This concludes our discussion of the age-less model, which was used to calculate individual parameters in an unbiased way.

To adequately assess the age trajectories of fitted parameters, we employed a fitting technique based on hierarchical Bayesian model fitting (Katahira, 2016; M. D. Lee, 2011), which avoids biases that arise when comparing parameters between participants that have been fitted using maximum-likelihood (van den Bos et al., 2017), and allows to test specific hypotheses about parameter trajectories by explicitly modeling these trajectories within the fitting framework: We conducted a separate “age-based” model, in which model parameters were allowed to depend on participants’ age (Fig. 3B). Estimating age effects directly within the computational model allowed us to estimate group-level effects in an unbiased way, whereas flat (hierarchical) models that estimate parameters but not age effects would underestimate (overestimate) group-level effects, respectively (Boehm et al., 2018). The age-based model was exclusively used to statistically assess parameter age trajectories because individual parameters would be biased by the inclusion of age in the model.

In the age-based model, each parameter *θ* of each participant *j* was sampled from a Normal distribution around an age-based regression line (Fig. 3B):

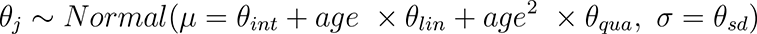

Each parameter’s intercept *θ_int_*, linear change with age *θ_lin_*, quadratic change with age *θ_qua_*, and standard deviation *θ_sd_* were sampled from prior distributions of the form specified in suppl. Table 5.

#### 4.5.4. Correlations between Model Parameters (*Fig. 5B*)

We used Spearman correlation because parameters followed different, not necessarily normal, distributions. Employing Pearson correlation led to similar results. p-values were corrected for multiple comparisons using the Bonferroni method.

#### 4.5.5. Principal Component Analysis (PCA)

To extract general cognitive components from model parameters, we ran a PCA on all fitted parameters (8 per participant). PCA can be understood as a method that rotates the initial coordinate system of a dataset (in our case, 8 axes corresponding to the 8 parameters), such that the first axis is aligned with the dimension of largest variation in the dataset (first principle component; PC1), the second axis with the dimension of second-largest variance (PC2), while being orthogonal to the first, and so on. In this way, all resulting PCs are orthogonal to each other, and explain subsequently less variance in the original dataset. We conducted a PCA after centering and scaling (z-scoring) the data, using R (RCoreTeam, 2016).

To assess PC age effects, we ran similar regression models as for behavioral measures, predicting PCs from z-scored age (linear), z-scored age (quadratic), and sex. When significant, effects were noted in Fig. 5E. For PC2 and PC4, we also conducted post-hoc t-tests, correcting for multiple comparison using the Bonferroni method (suppl. Table 15).

## 5. Acknowledgments

Numerous people contributed to this research: Amy Zou, Lance Kriegsfeld, Celia Ford, Jennifer Pfeifer, Megan Johnson, Gautam Agarwal, Liyu Xia, Vy Pham, Rachel Arsenault, Josephine Christon, Shoshana Edelman, Lucy Eletel, Neta Gotlieb, Haley Keglovits, Julie Liu, Justin Morillo, Nithya Rajakumar, Nick Spence, Tanya Smith, Benjamin Tang, Talia Welte, and Lucy Whitmore. We are also grateful to our participants and their families. The work was funded by National Science Foundation SL-CN grant 1640885 to RD, AGEC, and LW.

## 6. Supplemental Material

### 6.1. Supplemental Introduction

#### 6.1.1. Overview of Previous Reversal-Learning Studies in Adolescents

We know of three other groups that have investigated the development of reversal learning before. Table 3 shows the methods used in these studies, and Table 4 summarizes the main findings. It is of note that others have investigated developing populations on reversal tasks as well, but either age effects were not recorded (e.g., due to a focus on clinical questions; Adleman et al., 2011; Boehme et al., 2017; Dickstein, Finger, Brotman, et al., 2010; Dickstein, Finger, Skup, et al., 2010; Finger et al., 2008; Harms et al., 2018), or participants were younger and studies did not include adolescents (e.g., Minto de Sousa et al., 2015).

**Table 3:** Overview of studies that have used reversal tasks in human adolescents and investigated age effects. This table details participant samples, task designs, and RL modeling methods.

| Study | Participant age | Task | RL model | RL model quality |
| --- | --- | --- | --- | --- |
| Javadi et al., 2014 | 14-15 (n=260)<br>20-39 (n=29) | Select one stimulus on each trial<br>Correct: 70% reward, 30% punishment<br>Incorrect: 40% reward, 60% punishment<br>Reversal: 25% per trial after $\geq 4$ correct | Adaptive $\alpha$<br>(3 parameters) | No model comparison<br>No model validation |
| Hauser et al., 2015 | 12-16 (n=19)<br>20-29 (n=17) | Select one stimulus on each trial<br>Correct: 80% reward, 20% punishment<br>Incorrect: 20% reward, 80% punishment<br>Reversal: 6-10 trials after 3 consec. correct | [Positive vs negative] and<br>[factual vs count.-fact.] learn. rates | Model comparison (3 models)<br>No model validation |
| van der Schaaf et al., 2011 | 10-11 (n=15)<br>13-14 (n=15)<br>16-17 (n=15)<br>20-25 (n=16) | Predict outcome of highlighted stimulus<br>Correct: 100% reward, 0% punishment<br>Incorrect: 0% reward, 100% punishment<br>Reversal: after 4-6 consecutive correct | No computational model |  |
| Ours | 8-10 (n=41)<br>10-13 (n=46)<br>13-15 (n=45)<br>15-17 (n=47)<br>18-26 (n=57)<br>25-30 (n=55) | Select one stimulus on each trial<br>Correct: 75% reward, 25% punishment<br>Incorrect: 0% reward, 100% punishment<br>Reversal: After 7-15 rewards | [Positive vs negative] and<br>[factual vs count.-fact.] learn. rates;<br>persistence | Extensive model comparison<br>(7 RL & 16 BI models)<br>Extensive model validation |

**Table 4:** Overview of the results of the studies in suppl. Table 3. We focus on age differences in overall performance, the number of reversals (another performance measure), and RL model parameters. Note that differences between studies need to be interpreted carefully because task design, participant samples, and computational models differed between studies, as shown in suppl. Table 3.

| Study | Performance | Number of reversals | RL model results |
| --- | --- | --- | --- |
| Javadi et al., 2014 | No age difference | More in adults | $\log(\gamma)$ lower in adolescents<br>Larger RPEs in adolescents after<br>correct responses but negative feedback |
| Hauser et al., 2015 | No age difference | No age difference | $\alpha_-$ *factual* higher in adolescents |
| van der Schaaf et al., 2011 | Linear increase non-reversal trials<br>(asymptote in adolescence)<br>Inverse U-shape reversal trials<br>(max in adolescence) | Linear increase with age | No model |
| Ours | Inverse U-shape asymptotic trials<br>(max in mid-adolescence)<br>Inverse U-shape reversal trials<br>(max in adolescence) | NA | $p$ increases with age, asymptotes in late adolescence<br>$\beta$ increases with age, asymptotes in late adolescence<br>$\alpha_-$ U-shape, lowest in mid-adolescence<br>$\alpha_+$ step function, larger in adults |

### 6.2. Supplemental Methods

#### Quantile Age Bins

For some analyses, we split participants into quantiles based on age. This data binning led to samples of adequate sizes for summary statistics, while re-balancing group sizes after participant exclusion (see section 4.1). For participants below 18 years, quantiles were created by first separating males and females. For each sex, we then determined the cut-off ages that created the most balanced groups in terms of participant numbers, and recombined males and females to ensure even proportions of males and females in each age bin. For adult participants, we split the sample at 25 years of age.

#### 6.2.1. Comparing the Effectiveness of Hierarchical Bayesian Model Fitting versus Maximum-Likelihood Fitting on the Current Task

All model fits are relative: When model A fits data better than model B, there is no guarantee that model A fits the data “well”. Both models could fit the data poorly, with model A fitting just slightly better than model B. To ensure that our models fit well, we therefore validated parameter fitting and model comparison by first simulating and then recovering parameters from each model (Palminteri et al., 2017; Wilson and Collins, 2019). An identifiable model will recover the simulated parameters well during fitting, whereas an unidentifiable model will not. We also compared the results of maximum likelihood and hierarchical Bayesian model fitting using this procedure.

Figure 6A shows the well-established finding that hierarchical Bayesian model fitting outperforms the maximum likelihood method (Katahira, 2016): Both BF and RL model parameters were recovered well when using hierarchical Bayesian model fitting (age-free model), but not when using maximum likelihood. Furthermore, hierarchical Bayesian model fitting led to more consistent estimates of parameters *β* and *p* between both models (suppl. Fig. 6B), showing that this method was especially suited for our dual-model approach. These results lend credence to the superior fit that can be achieved using Hierarchical Bayesian methods, and to the precision with which model parameter can be estimated.

**Figure 6:**
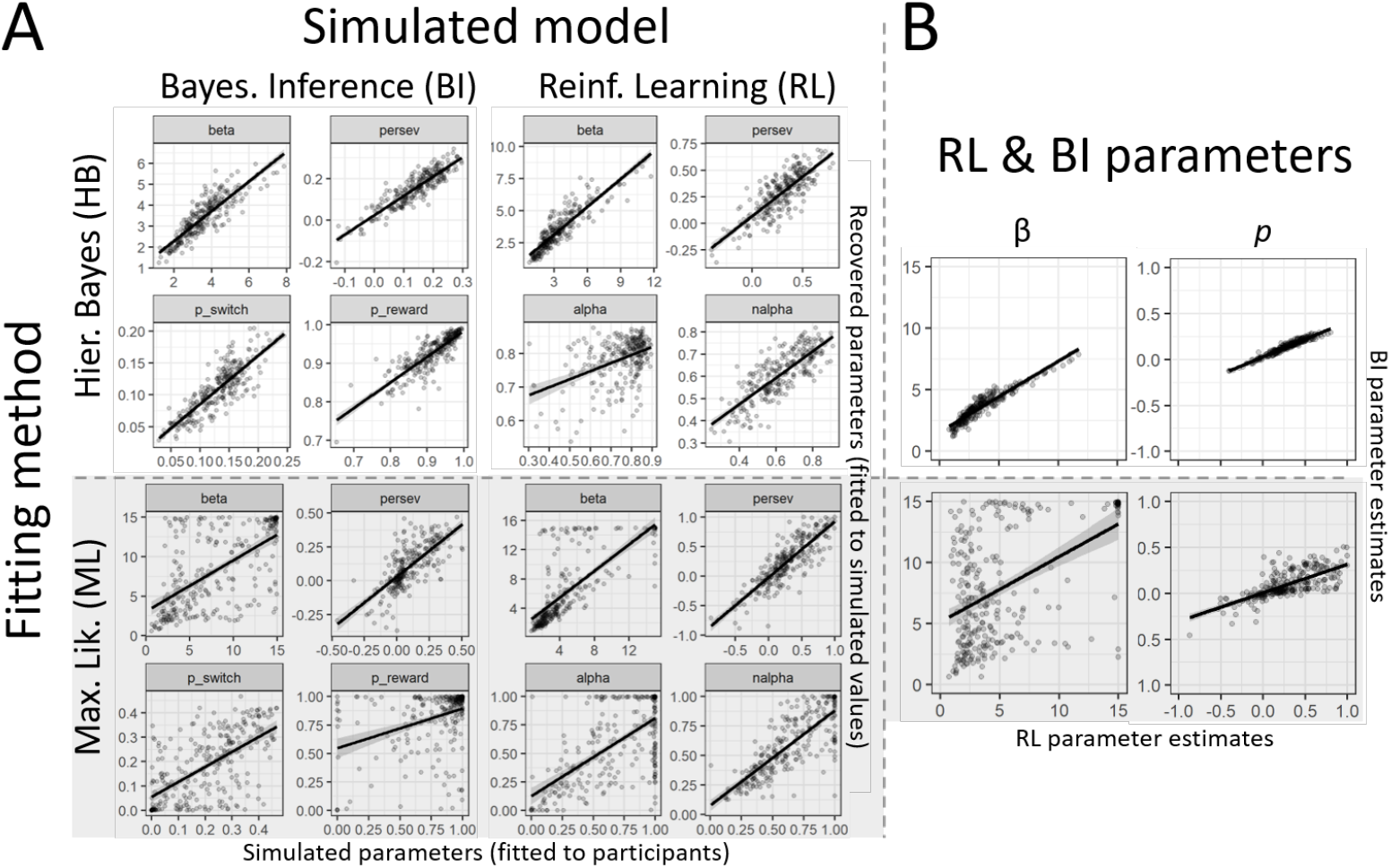
Model validation using hierarchical Bayesian model fitting (top, unshaded) and Maximum likelihood fitting (bottom, shaded). A) Simulate-and-recover procedure. The x-axes of all graphs show the parameter values of simulated datasets; the y-axes show the recovered parameters obtained by fitting these datasets using the same models. Recovered parameters should be as close to the simulated ones as possible, i.e., lie on the identity line. Black lines and shaded areas indicate best-fit regression lines. The left half presents simulate-and-recover results for the BI model, the right for the RL model. The top half shows the results of hierarchical Bayesian model fitting (our method), the bottom of the maximum likelihood method (standard). B) Consistency in the estimation of parameters *β* and *p*. Human data was fit using RL and BI models to compare the estimates of *β* (left row) and *p* (right row) between models. When both—independent—models lead to the same estimates, dots lie on the identity line. This was indeed the case for hierarchical Bayesian fitting (top row), but not for maximum likelihood fitting (bottom row).

#### 6.2.2. Hierarchical Bayesian Model Fitting

Hierarchical Bayesian model fitting requires the choice of the shapes of the prior distributions from which individuals’ parameters are drawn, and in some cases the choice of the distributions and parameters from which the parameters of the prior distributions are drawn. These choices can potentially influence fitting results; we chose non-informative prior distributions to limit the effect of these choices on our results. Table 5 shows the chosen distribution shapes and parameters.

**Table 5:**
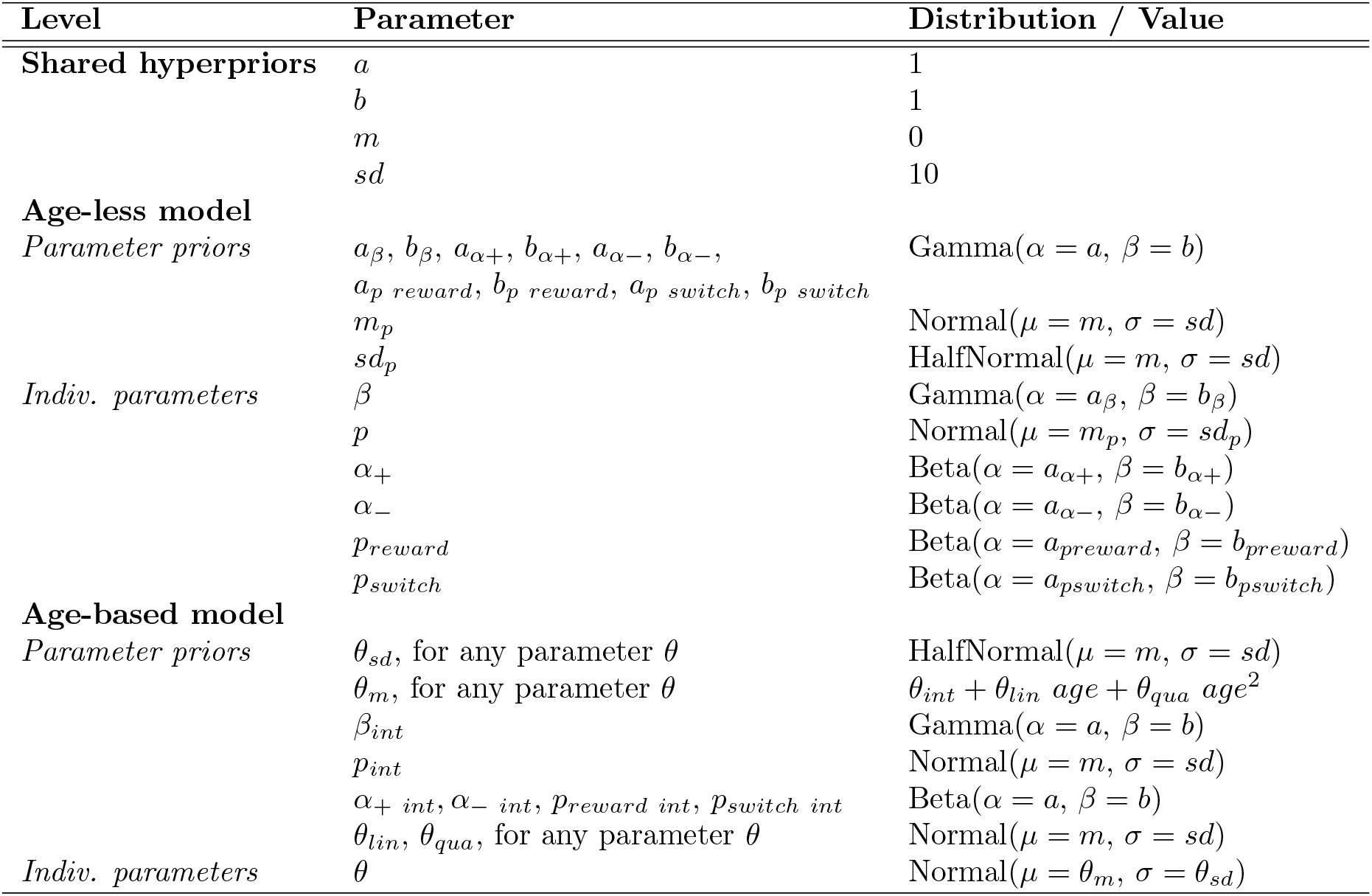
Priors and hyper-priors used in hierarchical Bayesian model fitting (Fig. 7B), chosen to be uninformative.

| Level | Parameter | Distribution / Value |
| --- | --- | --- |
| **Shared hyperpriors** | $a$ | 1 |
| | $b$ | 1 |
| | $m$ | 0 |
| | $sd$ | 10 |
| **Age-less model** |  |  |
| *Parameter priors* | $a_\beta, b_\beta, a_{\alpha+}, b_{\alpha+}, a_{\alpha-}, b_{\alpha-},$ | Gamma($\alpha = a, \beta = b$) |
| | $a_p \text{ reward}, b_p \text{ reward}, a_p \text{ switch}, b_p \text{ switch}$ | |
| | $m_p$ | Normal($\mu = m, \sigma = sd$) |
| | $sd_p$ | HalfNormal($\mu = m, \sigma = sd$) |
| *Indiv. parameters* | $\beta$ | Gamma($\alpha = a_\beta, \beta = b_\beta$) |
| | $p$ | Normal($\mu = m_p, \sigma = sd_p$) |
| | $\alpha_+$ | Beta($\alpha = a_{\alpha+}, \beta = b_{\alpha+}$) |
| | $\alpha_-$ | Beta($\alpha = a_{\alpha-}, \beta = b_{\alpha-}$) |
| | $p_{\text{reward}}$ | Beta($\alpha = a_{p_{\text{reward}}}, \beta = b_{p_{\text{reward}}}$) |
| | $p_{\text{switch}}$ | Beta($\alpha = a_{p_{\text{switch}}}, \beta = b_{p_{\text{switch}}}$) |
| **Age-based model** |  |  |
| *Parameter priors* | $\theta_{sd}$, for any parameter $\theta$ | HalfNormal($\mu = m, \sigma = sd$) |
| | $\theta_m$, for any parameter $\theta$ | $\theta_{int} + \theta_{lin} \text{ age} + \theta_{qua} \text{ age}^2$ |
| | $\beta_{int}$ | Gamma($\alpha = a, \beta = b$) |
| | $p_{int}$ | Normal($\mu = m, \sigma = sd$) |
| | $\alpha_+ \text{ int}, \alpha_- \text{ int}, p_{\text{reward int}}, p_{\text{switch int}}$ | Beta($\alpha = a, \beta = b$) |
| | $\theta_{lin}, \theta_{qua}$, for any parameter $\theta$ | Normal($\mu = m, \sigma = sd$) |
| *Indiv. parameters* | $\theta$ | Normal($\mu = \theta_m, \sigma = \theta_{sd}$) |

##### Prior Distributions for Individual Parameters

As shown in the table, in the age-based model (see section 4.5.3 for the differentiation between the age-based and age-free models), individuals’ parameters were drawn from a Normal distribution around a parameter-specific, continuously age-dependent mean *θ_m_*, with parameter-specific standard deviation *θ_sd_*.

In the age-free model, on the other hand, individuals’ parameters were drawn from parameter-specific group-level prior distributions. The shapes of these distributions were based on allowed parameter ranges (e.g., Gamma distribution for parameters with range [0, ∞], Beta distribution for parameters with range [0, 1]). The same prior distribution was used for all individuals, i.e., no age information was present in the age-free model. The distributions of individuals’ parameters were themselves parameterized by prior parameters.

##### Hyper-Prior Distributions of Prior Distribution Parameters

As further shown in the table, in the age-based model, prior parameter *θ_sd_* was distributed according to a HalfNormal (Normal, truncated at 0 to leave only support *>* 0), and parameterized by hyper-parameter *sd* = 10 to allow for a wide, non-informative shape. Group-level prior *θ_m_* was defined as an age-based regression function, parameterized by *θ_int_*, *θ_lin_*, and *θ_qua_* for each parameter *θ*. The prior on the intercept *θ_int_* of each parameter in the age-based model had the same shape as the group-level prior distribution in the age-free model, and was parameterized by the same hyper-priors.

In the age-less model, prior parameters parameterized the distributions of individual model parameters.

It is important to verify the convergence of the Markov-Chain MonteCarlo (MCMC) chains that are used in hierarchical Bayesian model fitting to approximate the intractable posterior distributions over model parameters given a dataset *p*(*θ data*) (see section 4.5.3). To this aim, we calculated the Markov-Chain error, effective sample size, and the R-hat statistic (suppl. Table 6), using the functions provided by the PyMC3 toolbox (Salvatier et al., 2016).

**Table 6:** Convergence of MCMC chains used in hierarchical Bayesian model fitting. We report the Markov-Chain error, effective sample size (*n*), and the R-hat statistic (*R*^^^), showing averages and ranges (min and max over all model parameters) for both winning models.

| Model | | MC error | Effective $n$ | $\hat{R}$ |
| --- | --- | --- | --- | --- |
| **4-param. RL** | mean | < 0.001 | 2,517 | 1.001 |
|  | range | [< 0.001; 0.002] | [155; 4,261] | [1.000; 1.015] |
| **4-param. BI** | mean | 0.002 | 816 | 1.001 |
|  | range | [< 0.001; 0.01] | [281; 1,576] | [1.000; 1.004] |

One of our main questions in this research was whether model parameter changed with age. We used hierarchical Bayesian model fitting to address this question, given the possibility to assess age-related differences in computational model parameters in an unbiased way using this method (see section 4.5.3). In order to estimate individual (and group-level) parameters in hierarchical Bayesian model fitting, obtained MCMC samples are averaged; to test particular parameter hypotheses (e.g., a parameter is greater than 0), the proportion of samples is calculated in which the hypothesis is true, and this proportion can be compared to a pre-determined p-value to assess significance. Following this procedure, we determined whether the parameters in the age-based model that controlled the effect of age on model parameters showed significant differences from 0. Table 13 shows the results, revealing significant linear and quadratic effects for some parameters.

### 6.3. Supplemental Results

#### 6.3.1. Additional Behavioral Measures

We analyzed participant behavior in more detail than presented in the main text. For example, we completed the assessment of performance by analyzing the number of points won by each participant suppl. Fig. 7A, E), we assessed flexibility by counting the trials it took participants between a task switch to complete a behavioral switch (lower is faster; suppl. Fig. 7B, F). We also assessed the effects of positive (suppl. Fig. 7D, H) and negative ((suppl. Fig. 7C, G) outcomes on subsequent actions, using the regression analysis described in section 4.4, whose statistics are reported in Table 7 below. Each behavior showed interesting age trajectories.

**Figure 7:**
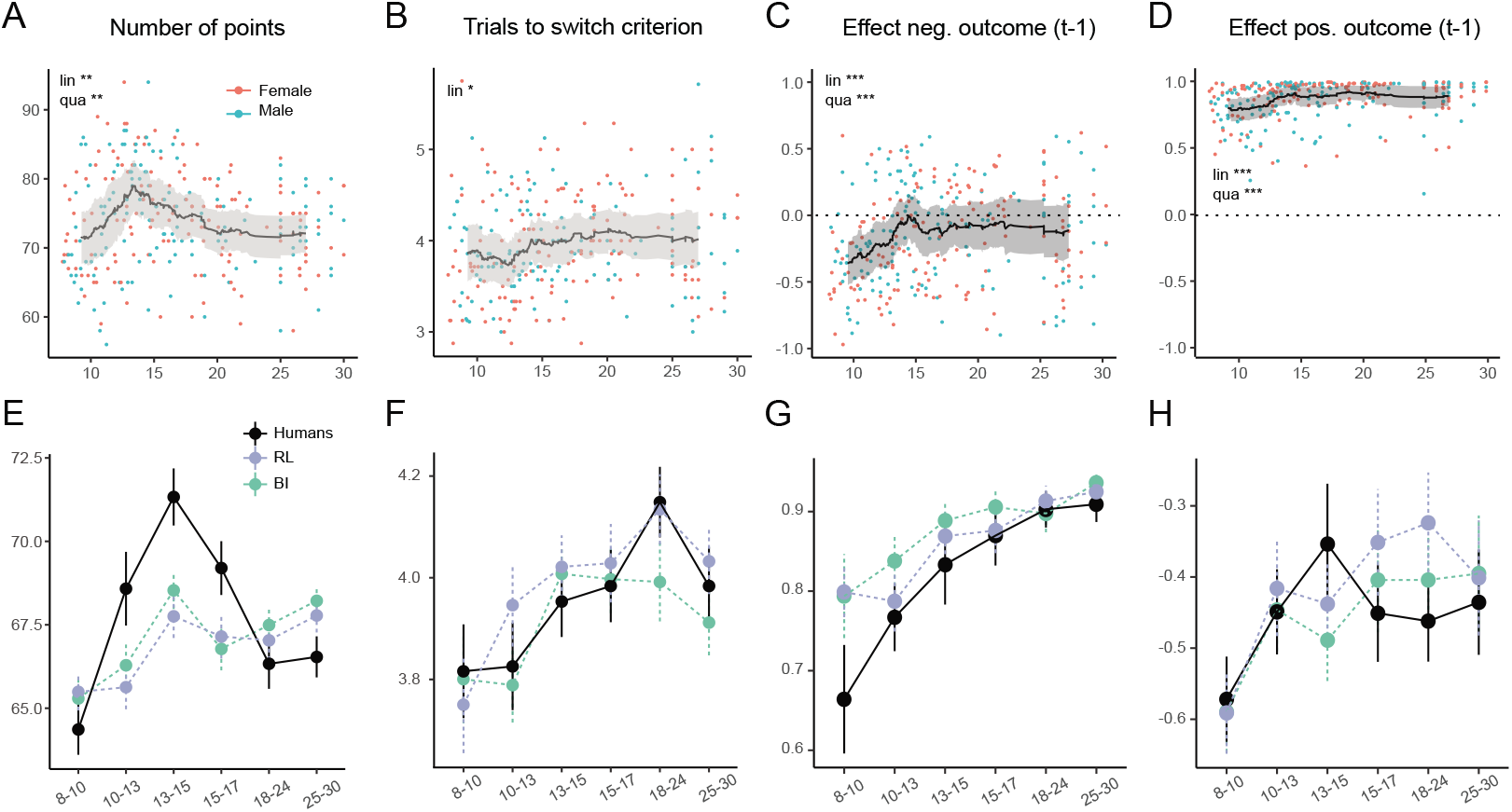
Human behavior (A-D) and model validation (E-H) for additional behavioral measures. (A, E) Number of points won by each participant. (A) Each dot represents one participant, colors denote sex; the lines shows the rolling average, shades the standard error, highlighting the performance peak in mid- to late adolescence. (E) Number of points, averaged within age groups, showing human as well as model behavior for validation. (B, F) Number of trials after task switch until participants reached performance criterion (2 correct responses). (C, D, G, H) Effect of previous negative (C, G) or positive (D, H) outcomes on participants’ choices. “*t* − 1”: The assessed outcome occurred 1 trial before choice, i.e., delay *i* = 1. Regression weights were tanh transformed for visualization. The youngest age groups showed the lowest overall and asymptotic accuracy (main text Fig. 3C, F) and were most likely to switch after a single negative outcome (main text Fig. 3E, suppl. Fig. 15B, middle). This explains why they were also fastest at switching (this Figure, parts B and F).

**Table 7:** Logistic mixed-effect regression, predicting future actions from past actions and outcomes. The number of predictors (*i ≤* 8) was chosen as to provide the best model fit: *AIC_i≤_*_3_: 31.046; *AIC_i≤_*_4_: 31.013; *AIC_i≤_*_5_: 31.001; *AIC_i≤_*_6_: 30.981; *AIC_i≤_*_7_: 30.963; *AIC_i≤_*_8_**: 30.962**; *AIC_i≤_*_9_: 30.966; *AIC_i≤_*_10_: 30.964.

| Predictor | delay $i$ | $\beta$ | $z$ | $p$ | Sig. |
| --- | --- | --- | --- | --- | --- |
| **Intercept** |  | -0.01 | -0.74 | 0.46 |  |
| **Main effects** |  |  |  |  |  |
| Age (lin.) |  | -0.13 | -1.40 | 0.16 |  |
| Age (qua.) |  | 0.12 | 1.30 | 0.19 |  |
| Pos. outcome | 1 | 2.19 | 68.09 | < 0.001 | *** |
|  | 2 | 0.84 | 27.36 | < 0.001 | *** |
|  | 3 | 0.24 | 7.87 | < 0.001 | *** |
|  | 4 | 0.13 | 4.30 | < 0.001 | *** |
|  | 5 | -0.017 | -0.54 | 0.58725 |  |
|  | 6 | -0.017 | -0.56 | 0.57548 |  |
|  | 7 | -0.0035 | -0.12 | 0.90613 |  |
|  | 8 | -0.077 | -2.77 | 0.0057 | ** |
| Neg. outcome | 1 | -0.73 | -37.09 | < 0.001 | *** |
|  | 2 | -0.24 | -10.64 | < 0.001 | *** |
|  | 3 | 0.0055 | 0.22 | 0.82278 |  |
|  | 4 | 0.13 | 5.39 | < 0.001 | *** |
|  | 5 | 0.12 | 4.87 | < 0.001 | *** |
|  | 6 | 0.12 | 4.73 | < 0.001 | *** |
|  | 7 | 0.13 | 5.32 | < 0.001 | *** |
|  | 8 | 0.016 | 0.71 | 0.47857 |  |
| **Interaction age (lin.)** |  |  |  |  |  |
| Pos. outcome | 1 | 0.90 | 4.50 | < 0.001 | *** |
|  | 2 | 0.84 | 4.19 | < 0.001 | *** |
|  | 3 | 0.50 | 2.52 | 0.012 | * |
|  | 4 | -0.069 | -0.35 | 0.73 |  |
|  | 5 | 0.088 | 0.44 | 0.66 |  |
|  | 6 | -0.38 | -1.94 | 0.052 |  |
|  | 7 | -0.18 | -0.94 | 0.35 |  |
|  | 8 | -0.27 | -1.49 | 0.14 |  |
| Neg. outcome | 1 | 0.67 | 5.27 | < 0.001 | *** |
|  | 2 | -0.37 | -2.48 | 0.013 | * |
|  | 3 | 0.16 | 1.03 | 0.30 |  |
|  | 4 | -0.089 | -0.55 | 0.58 |  |
|  | 5 | 0.012 | 0.07 | 0.94 |  |
|  | 6 | 0.066 | 0.41 | 0.68 |  |
|  | 7 | 0.011 | 0.07 | 0.94 |  |
|  | 8 | -0.068 | -0.47 | 0.63 |  |
| **Interaction age (qua.)** |  |  |  |  |  |
| Pos. outcome | 1 | -0.64 | -3.14 | 0.0017 | ** |
|  | 2 | -0.89 | -4.41 | < 0.001 | *** |
|  | 3 | -0.38 | -1.90 | 0.057 |  |
|  | 4 | 0.0020 | 0.01 | 0.99 |  |
|  | 5 | -0.066 | -0.33 | 0.74 |  |
|  | 6 | 0.36 | 1.80 | 0.072 |  |
|  | 7 | 0.15 | 0.75 | 0.456 |  |
|  | 8 | 0.29 | 1.62 | 0.11 |  |
| Neg. outcome | 1 | -0.56 | -4.34 | < 0.001 | *** |
|  | 2 | 0.30 | 2.00 | 0.046 | * |
|  | 3 | -0.16 | -0.97 | 0.33 |  |
|  | 4 | 0.092 | 0.57 | 0.57 |  |
|  | 5 | -0.0070 | -0.04 | 0.97 |  |
|  | 6 | -0.092 | -0.57 | 0.57 |  |
|  | 7 | -0.057 | -0.35 | 0.72 |  |
|  | 8 | 0.064 | 0.44 | 0.66 |  |

#### 6.3.2. Comparing Behavioral Measures between Adolescents and Other Age Groups

In terms of which behavioral measures, and compared to which specific age groups, did adolescents perform better? For completeness, Table 8 reports the results of t-tests comparing the age bin of 13-to-15-year-olds to each other age group, in each performance measure. All tests were corrected for multiple comparisons using the Bonferroni method.

**Table 8:**
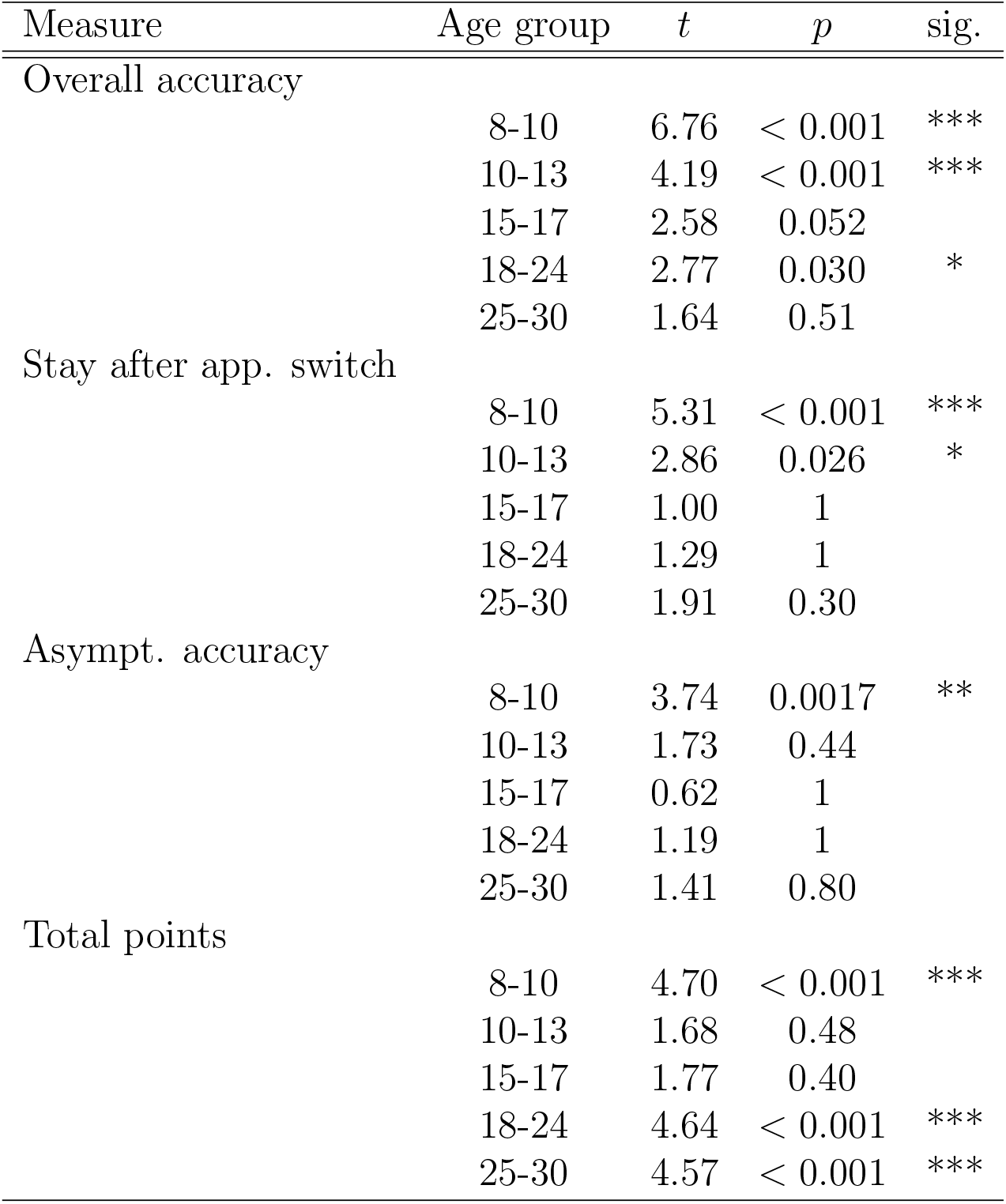
T-tests comparing participants in the 13-to-15-year-old age bin to all other age groups in terms of overall accuracy (Fig. 3C), stay after apparent switch (Fig. 3E), accuracy on asymptotic trials (Fig. 3F), and total points won (Fig. 2B). Each row shows the comparison of mid- to late adolescence to one other age group.

#### 6.3.3. An Effect of Puberty?

As mentioned in the Discussion, our results show age differences in the adaptation to stochastic and volatile environments, but do not identify a biological mechanism that underlies these differences. One possibility are puberty-related changes. To address this possibility, we asked participants aged 8-17 to complete the pubertal developmental scale (PDS), a questionnaire that determines pubertal status based on questions about physical development (Petersen et al., 1988), and to provide a 1.8 ml saliva sample, which was analyzed for testosterone levels as a marker of pubertal development, an hour after the start of the experiment and in-between tasks (for detailed methods, see Master et al., 2020). We then investigated how performance and model parameters changed with pubertal development, assessed using these two measures. We found qualitatively similar developmental patterns for puberty as for age (suppl. Fig. 9, 10, 11; suppl. Tables 10, 11), making it difficult to disentangle the effects of both because pubertal measures were highly correlated with age (suppl. Fig. 8). To investigate whether pubertal development had a unique effect after controlling for age, we also tested puberty effects within age bins, but failed to observe differences that were statistically significant (suppl. Fig. 12, 13, 14).

Nevertheless, some trends that emerged in the pubertal analyses, especially in pre-pubertal participants, deserve a more detailed investigation in future research, potentially employing longitudinal designs for enhanced experimental control (Kraemer et al., 2000).

**Figure 8:**
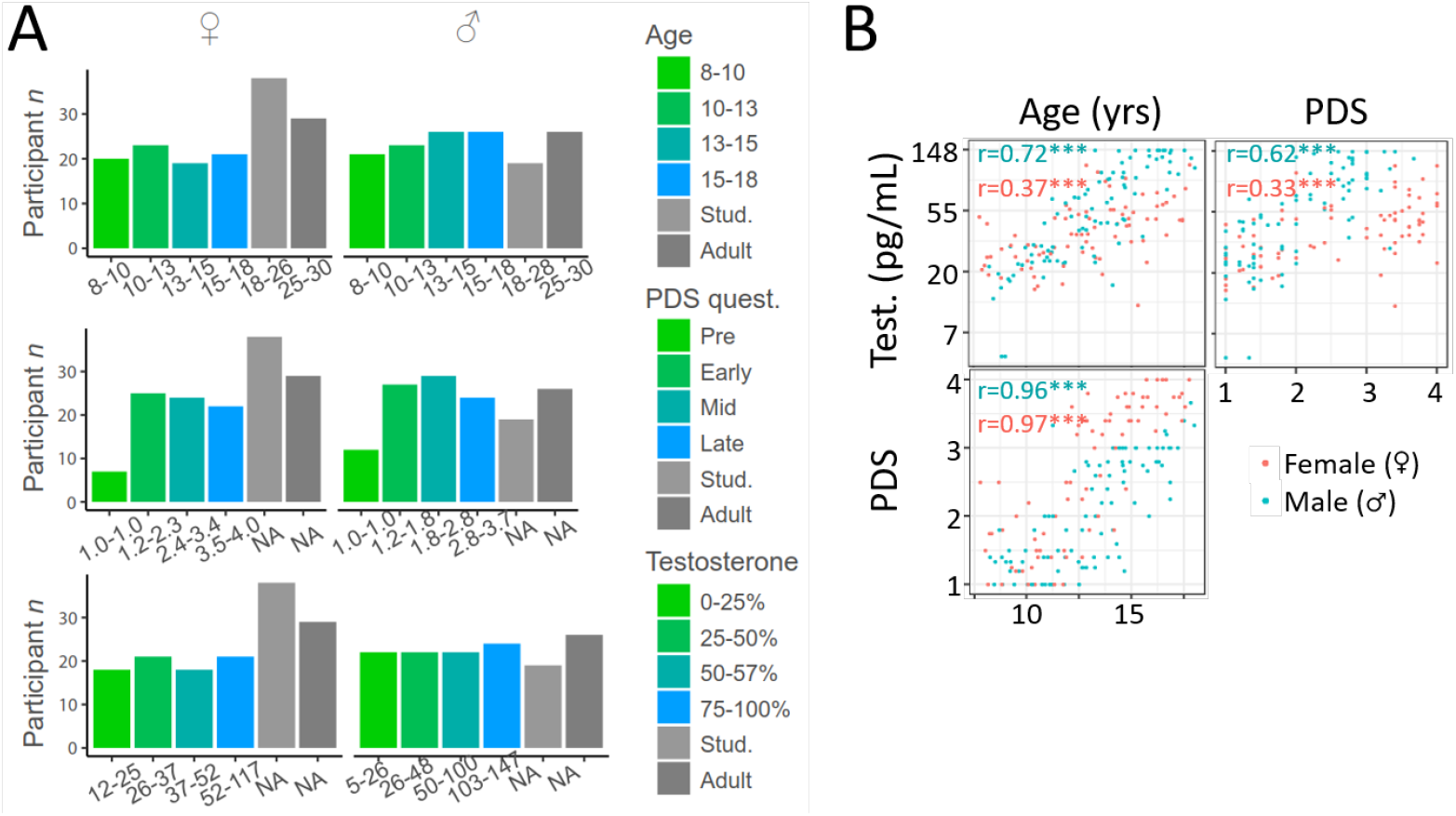
A) Participant numbers for each age bin (top), PDS score bin (middle), and Testosterone level bin (bottom). Pubertal measures were available for participants aged 8-17, and quantile bins were calculated in a similar way as for age, with one exception: For PDS scores, all participants with score 1 were classified as pre-pubertal, and the binning was only only conducted for remaining participants. Note that PDS and testosterone ranges differed substantially between sexes. B) Correlations between age, testosterone levels (Test.), and PDS questionnaire, for male and female participants aged 8-17. Stars refer to p-values, using the same convention as in main text figures. For both males and females, PDS scores and testosterone levels were highly correlated with age, as well as with each other, making it difficult to assess these three factors separately.

**Figure 9:**
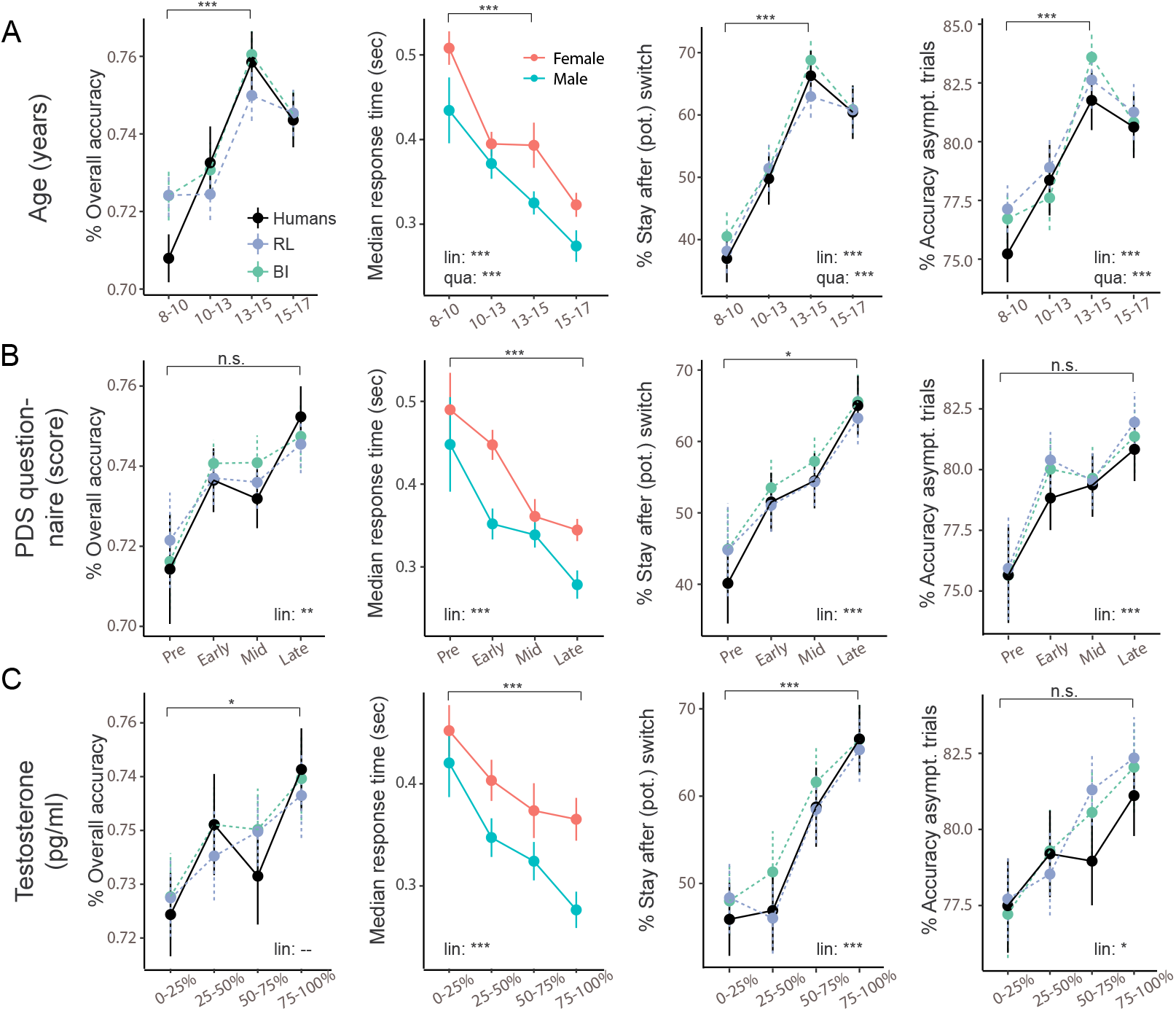
Behavior broken up by age (top row), PDS (middle row), and testosterone bins (bottom row). Significance bars and stars show the results of planned t-tests. A) Reproduced from main text Fig. 3. Planned t-tests compared 8-to-10-year-olds to 13-to-15-year-olds. B) Same data, but broken up by PDS bins. T-tests compared pre-pubertal to late-pubertal participants. C) Same data, broken up by testosterone bins. T-tests compared participants in the first quantile to participants in the fourth quantile. The figure shows that pubertal development (PDS, testosterone) was related to overall similar developmental patterns as age. The main difference lay in the bin of peak performance: Performance peaked in the third quantile based on age (13-15 years), but in the fourth quantiles based on PDS and testosterone.

**Figure 10:**
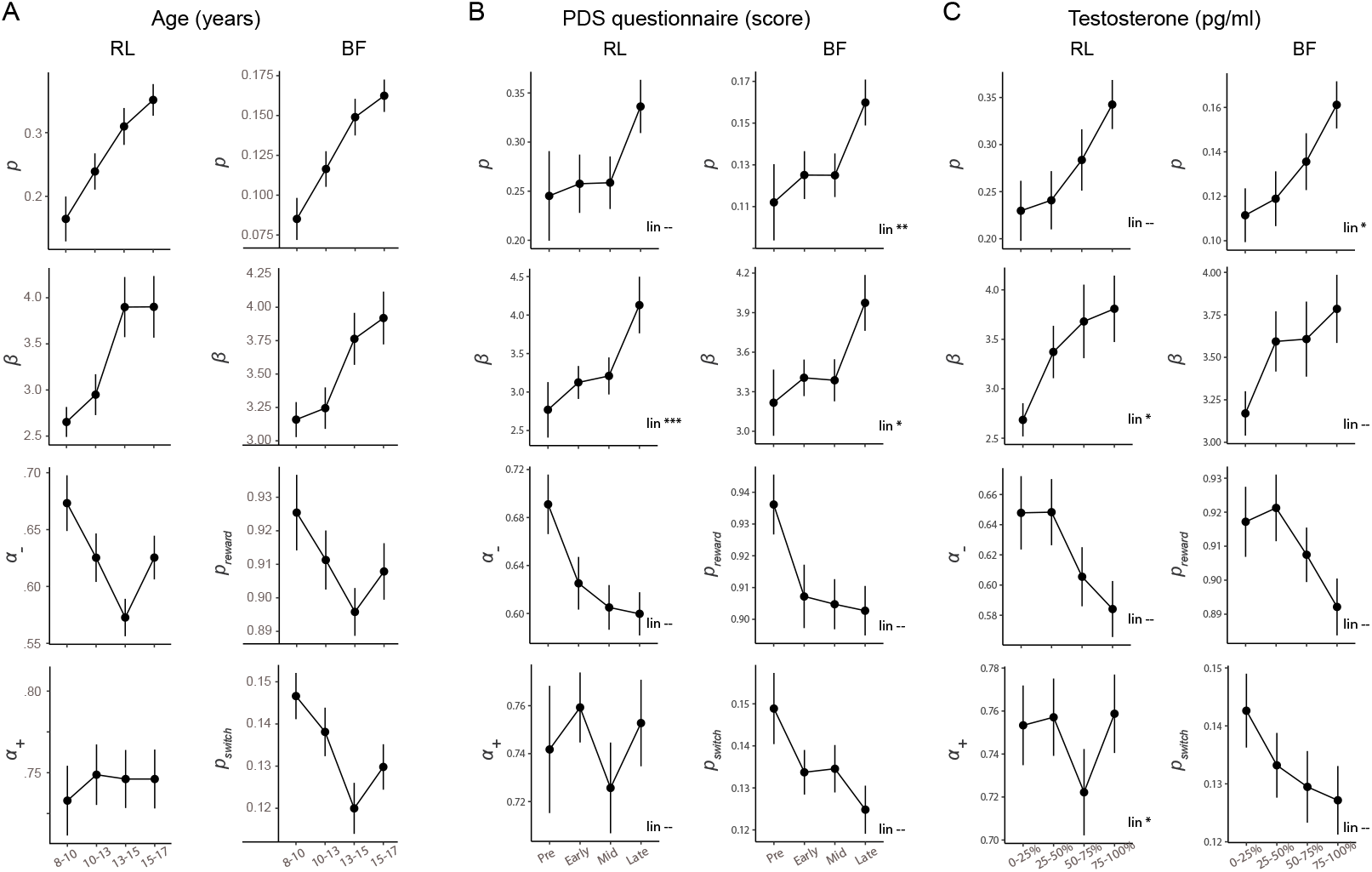
Model parameters broken up by age (A), PDS (B), and testosterone bins (C), showing that parameter trajectories varied slightly when analyzed through the lens of puberty compared to age. A) Reproduced from main text Fig. 4 after removing adult participants. B) Same data, broken up by PDS bins. Parameters *p* and *β* seem to show step functions between mid- and late puberty, as opposed to the gradual change with age (part A). Parameters *α_−_* and *p_reward_* seemed to show a drastic step at puberty onset (between “pre” and “early”), rather than the age-based U-shape. C) Same data, broken up by testosterone bins. Parameters *α_−_*, *p_reward_*, and *p_switch_* seemed to show U-shaped functions similar to age (elevated adult values are shown in main text Fig. 4), but minima occurred in the fourth rather than the third quantile. “lin.” indicates whether a linear effect of the measure of interest (PDS / testosterone) reached significance in a linear regression model.

**Figure 11:**
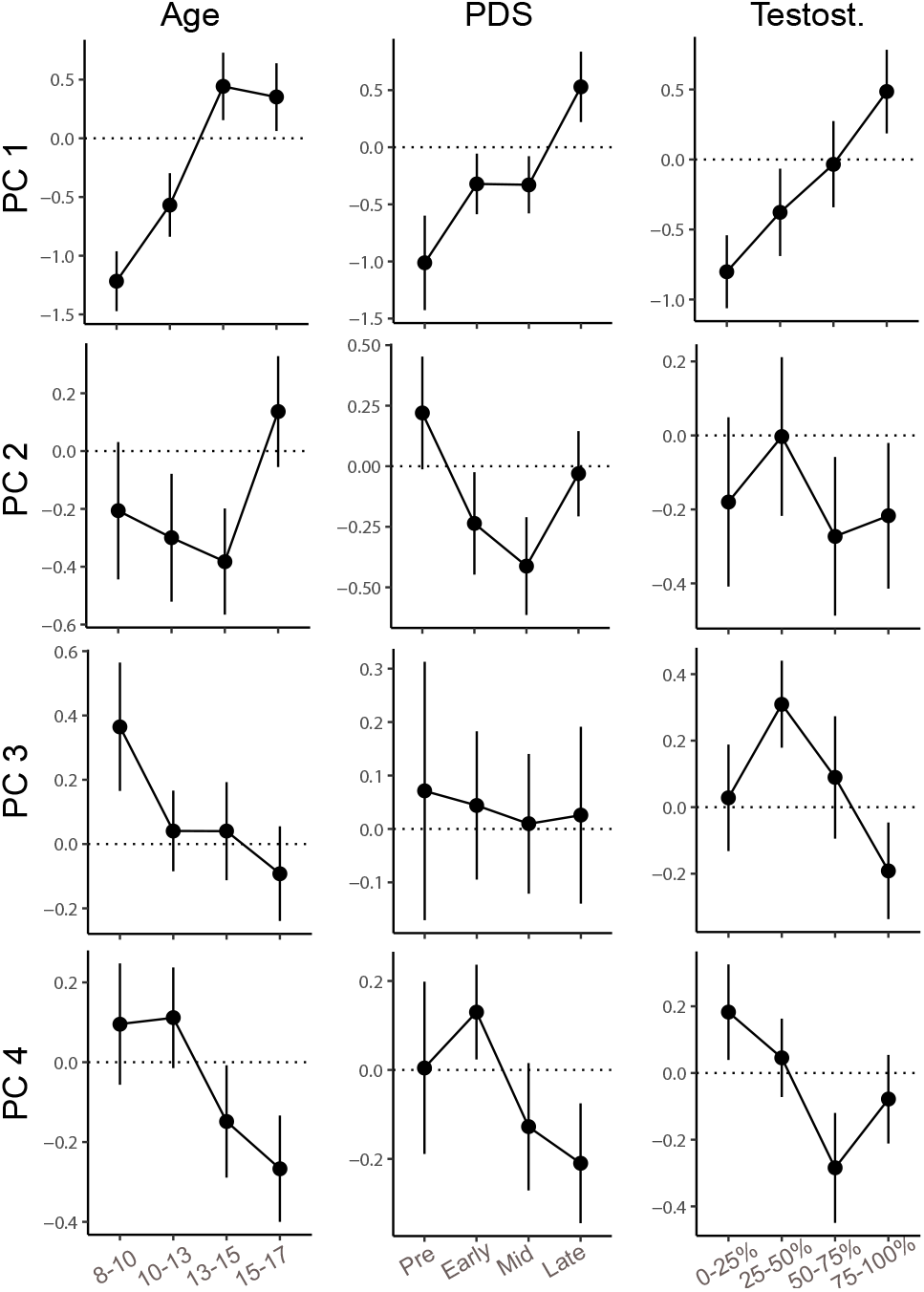
Model parameter PCs broken up by age (left), PDS (middle), and testosterone bins (right). Left: Reproduced from Fig. 5 after removing adult participants. Middle (right) row: same data, but broken up by PDS (testosterone) bins. This figure shows that in terms of parameter PCs, trajectories were relatively similar between pubertal measures and age. Slight differences included a more unique role of pre-pubertal participants, especially for PC2 in terms of PDS and PC3 for testosterone.

**Figure 12:**
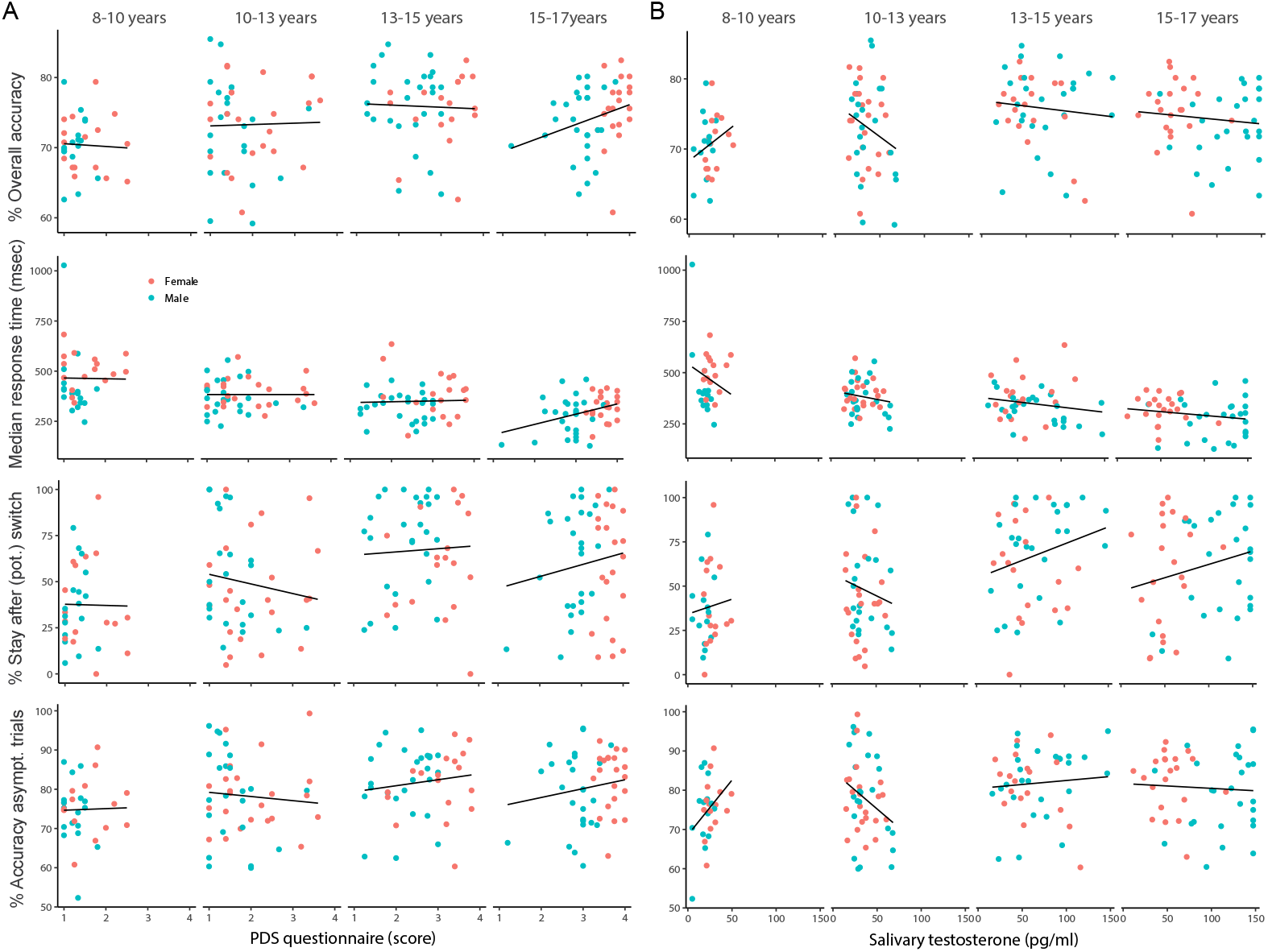
Effect of pubertal status on four performance measures, controlling for age. Each column shows one age group, colors denote sex. Pubertal status was determined by (A) PDS questionnaire, or (B) salivary testosterone. We sought to examine the effect of puberty after controlling for age. To this end, we investigated the continuous effects of puberty within each age bin, to eliminate—as much as possible—confounds with age (Master et al., 2020). A) In concordance with the finding that behavior peaked in the third age bin (13-15 years), but in the fourth PDS bin (75-100*^th^* percentile; suppl. Fig. 9), most performance measures increased qualitatively with respect to PDS in the third and fourth age bins (center-right and right-most column). Nevertheless, this pattern was difficult to interpret because pubertal status was heavily confounded with sex in the fourth age bin, such that girls scored higher on the PDS questionnaire than boys of the same age, a typical pattern that is caused by sex differences in pubertal maturation. It is therefore unclear whether the performance increase within the fourth age bin (right-most column) was driven by PDS scores or by sex. Stay after (pot.) switch trials showed a qualitative decrease with PDS score in 10-13 year olds, was constant in mid- to late adolescence, and showed a qualitative increase in 15-to-17-year-olds. This could indicate a weak U-shaped effect or might result from experimental noise. B) Same data, assessing age-controlled effects of testosterone on performance measures.

**Figure 13:**
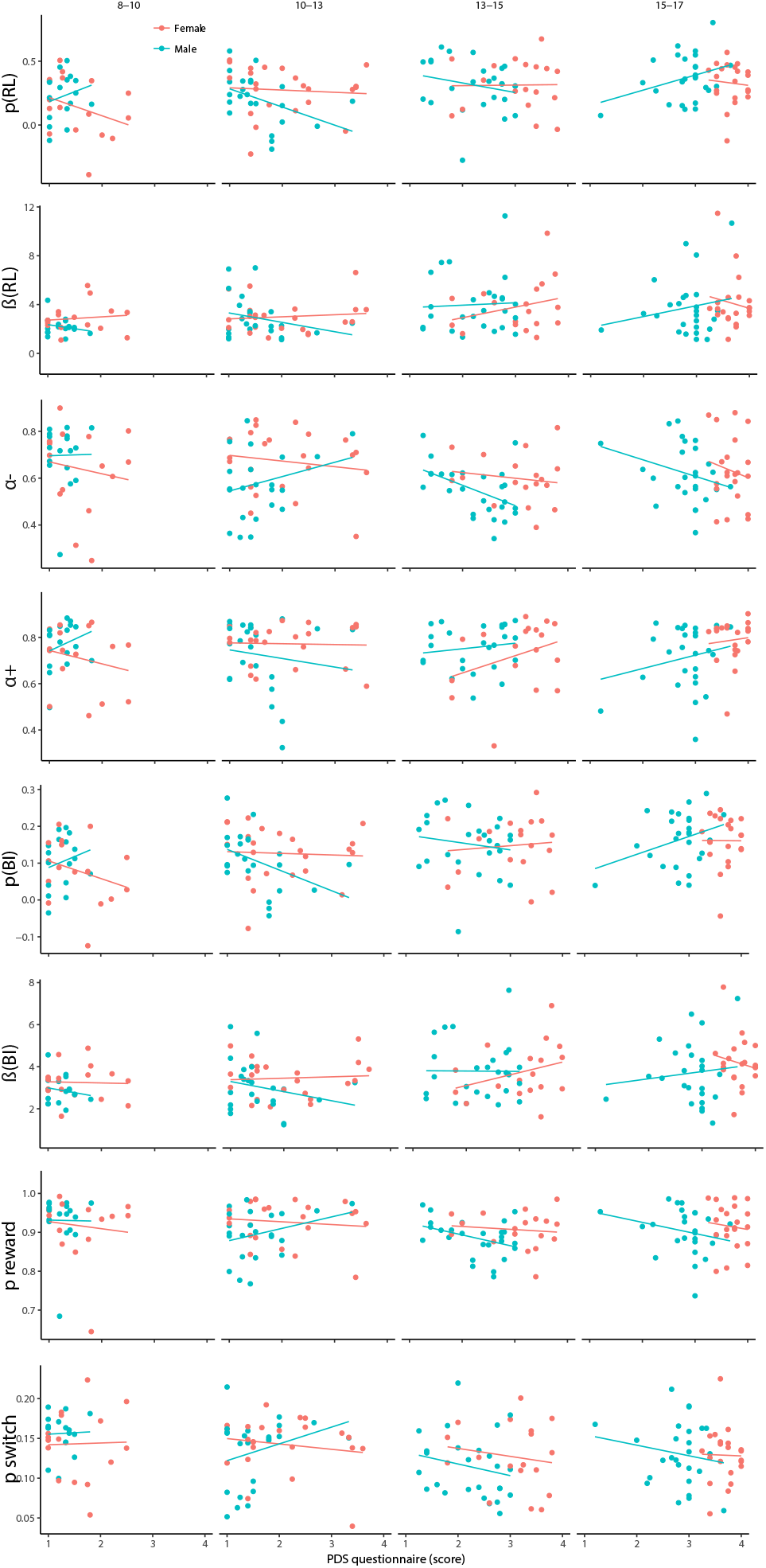
Effects of PDS scores on model parameters, controlling for age. Each column shows one age group, each row one parameter, and colors denote sex. Pubertal development did not show significant positive relationships with choice parameters *p* and *β*, which we might predict if pubertal development was a driving mechanism in growth for these parameters between ages 8-18 (see also suppl. Table 9 and suppl. Fig. 14). In terms of learning parameters, pubertal development also did not show significant negative relationships with *α_−_* and *α*_+_ (RL), or *p_reward_* and *p_switch_* (BI), which we might predict if pubertal onset was driving the decrease of these parameters between ages 8-15. If anything, we saw the opposite pattern in males: *α_−_*, *p_reward_*, and *p_switch_* showed a qualitatively positive relationship with PDS scores and testosterone (suppl. Fig. 14) in 10-to-13-year-olds, and a qualitatively negative relationship with PDS in mid- to late adolescence. Overwhelmingly, these relationships were not statistically significant (suppl. Table 9). Trend relationships within mid- to late adolescence included a marginal effect of PDS on *α*_+_ (suppl. Table 9). Note, however, that statistical tests were not corrected for multiple comparisons, making it possible that these results were observed by chance, and should thus be interpreted carefully. The cross-sectional design of our experiment may limit our ability to detect pubertal effects (Kraemer et al., 2000). It is possible that experiments with greater power, longitudinal studies, and studies of hormone manipulation may further inform these results.

**Figure 14:**
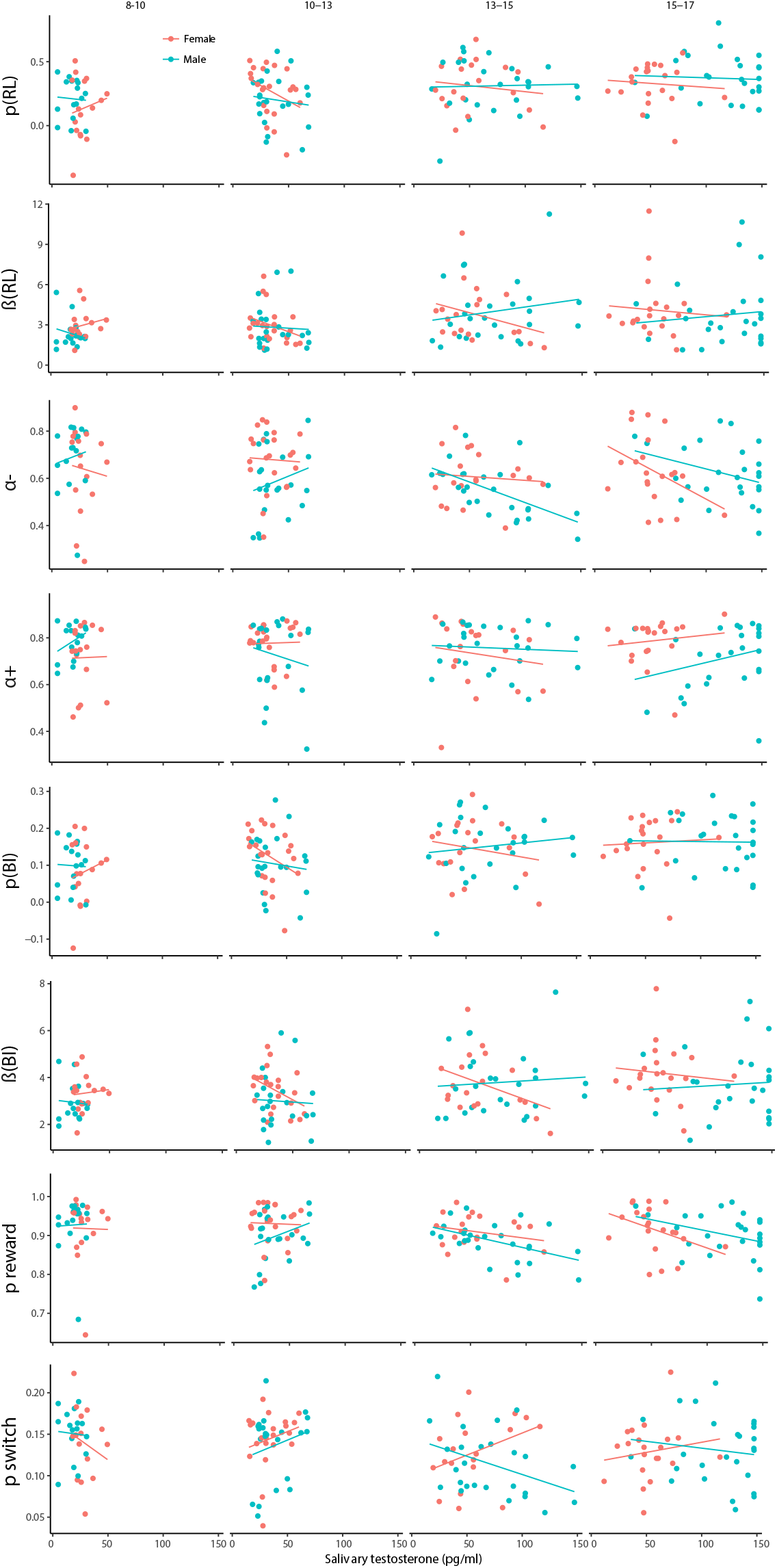
Effects of salivary testosterone levels on model parameters, controlling for age. Each column shows one age group, each row one parameter, and colors denote sex. Trend relationships within mid- to late adolescence included a marginal effect of sex on *p_switch_* in the testosterone model, and a significant interaction between sex and testosterone on *p_switch_* (suppl. Table 9). Note, however, that statistical tests were not corrected for multiple comparisons, making it possible that these results were observed by chance, and should thus be interpreted carefully.

**Table 9:** Statistics of regression models testing effects of puberty within the age bin 13-15 years. This bin was chosen because it contained participants across the full range of pubertal development.

| Outcome | Predictor | $\beta$ | $p$ | Sig. |
| --- | --- | --- | --- | --- |
| **Testosterone** |  |  |  |  |
| $p$ (RL) | Test. | -0.00096 | 0.57 | |
|  | Sex | 0.062 | 0.65 |  |
|  | Interaction | 0.0011 | 0.58 |  |
| $\beta$ (RL) | Test. | -0.022 | 0.23 | |
|  | Sex | 1.86 | 0.22 |  |
|  | Interaction | 0.034 | 0.13 |  |
| $\alpha_-$ | Test. | -0.00033 | 0.69 | |
|  | Sex | 0.047 | 0.48 |  |
|  | Interaction | 0.0014 | 0.16 |  |
| $\alpha_+$ | Test. | -0.00074 | 0.47 | |
|  | Sex | 0.0026 | 0.97 |  |
|  | Interaction | 0.00055 | 0.65 |  |
| $p$ (BF) | Test. | -0.00052 | 0.43 | |
|  | Sex | 0.045 | 0.40 |  |
|  | Interaction | 0.00083 | 0.30 |  |
| $\beta$ (BF) | Test. | -0.018 | 0.12 | |
|  | Sex | 1.12 | 0.21 |  |
|  | Interaction | 0.021 | 0.12 |  |
| $p_{reward}$ | Test. | -0.00038 | 0.31 | |
|  | Sex | 0.0012 | 0.97 |  |
|  | Interaction | 0.00027 | 0.54 |  |
| $p_{switch}$ | Test. | 0.00053 | 0.10 | |
|  | Sex | 0.047 | 0.078 | , |
|  | Interaction | 0.00097 | 0.015 | * |
| **PDS** |  |  |  |  |
| $p$ (RL) | PDS | 0.0044 | 0.95 | |
|  | Sex | 0.18 | 0.52 |  |
|  | Interaction | 0.079 | 0.43 |  |
| $\beta$ (RL) | PDS | 0.87 | 0.30 | |
|  | Sex | 2.37 | 0.45 |  |
|  | Interaction | 0.67 | 0.55 |  |
| $\alpha_-$ | PDS | -0.024 | 0.52 | |
|  | Sex | 0.071 | 0.61 |  |
|  | Interaction | 0.063 | 0.21 |  |
| $\alpha_+$ | PDS | 0.075 | 0.092 | , |
|  | Sex | 0.21 | 0.21 |  |
|  | Interaction | 0.051 | 0.39 |  |
| $p$ (BF) | PDS | 0.011 | 0.69 | |
|  | Sex | 0.084 | 0.45 |  |
|  | Interaction | 0.032 | 0.43 |  |
| $\beta$ (BF) | PDS | 0.62 | 0.21 | |
|  | Sex | 1.96 | 0.30 |  |
|  | Interaction | 0.64 | 0.34 |  |
| $p_{reward}$ | PDS | -0.0080 | 0.63 | |
|  | Sex | 0.023 | 0.72 |  |
|  | Interaction | 0.022 | 0.33 |  |
| $p_{switch}$ | PDS | -0.010 | 0.51 | |
|  | Sex | 0.010 | 0.86 |  |
|  | Interaction | 0.0057 | 0.82 |  |

**Table 10:**
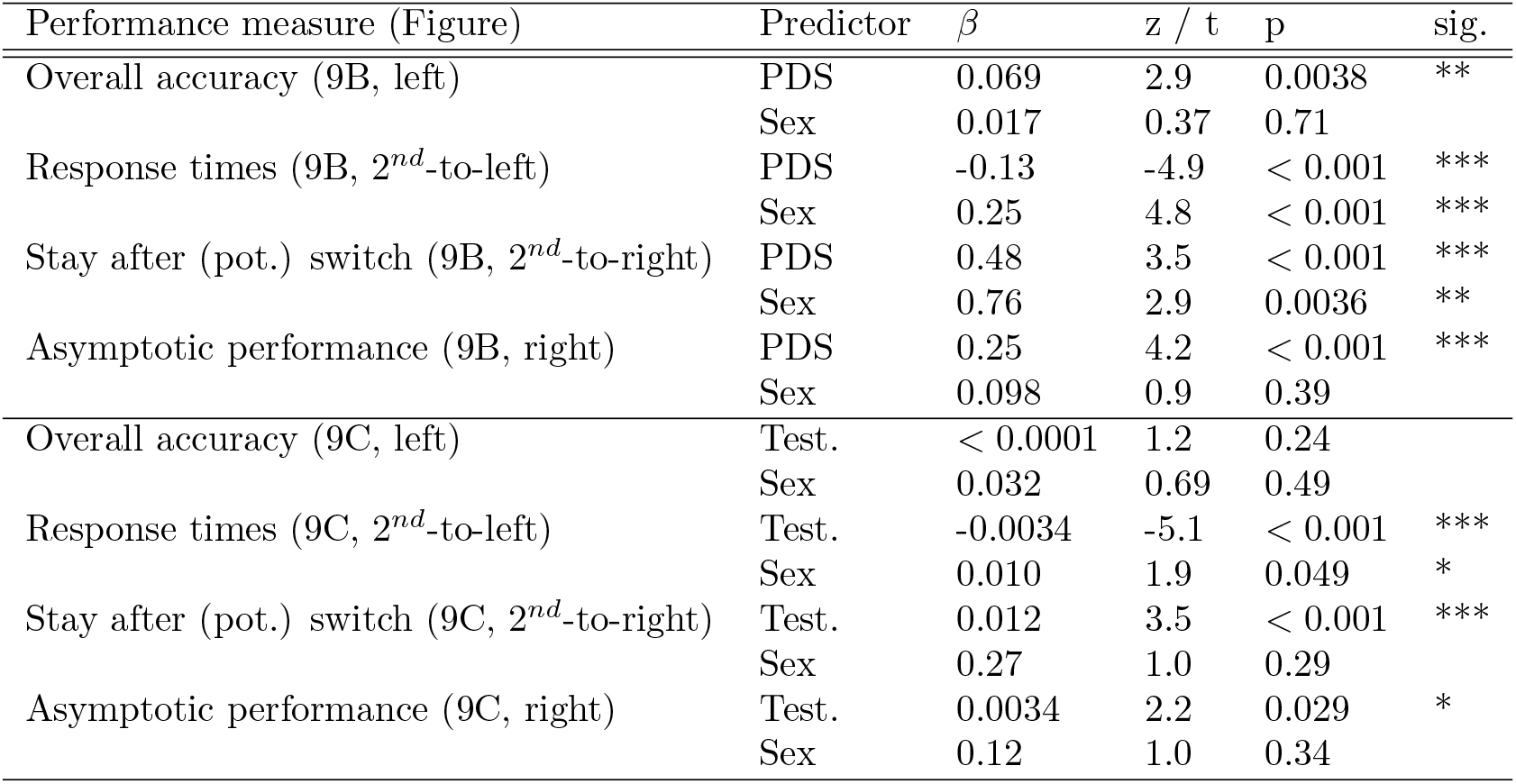
Statistics of mixed-effects regression models predicting performance measures from sex (male, female) and puberty measures (PDS questionnaire / salivary testosterone). Only participants aged 8-17 were included in this analyses because pubertal measures were only available for them. Overall accuracy, stay after potential (pot.) switch, and asymptotic performance were modeled using logistic regression, and z-scores are reported. Log-transformed response times on correct trials were modeled using linear regression, and t-values are reported. * *p < .*05; ** *p < .*01, *** *p < .*001. Within the age bins that contained participants across the entire range of pubertal status (10-13, 13-15, and 15-17 years), few significant effects of PDS (part A) or salivary testosterone levels (part B) were observed, possibly including some that occurred by chance.

| Performance measure (Figure) | Predictor | $\beta$ | z / t | p | sig. |
| --- | --- | --- | --- | --- | --- |
| Overall accuracy (9B, left) | PDS | 0.069 | 2.9 | 0.0038 | ** |
|  | Sex | 0.017 | 0.37 | 0.71 |  |
| Response times (9B, 2<sup>nd</sup>-to-left) | PDS | -0.13 | -4.9 | < 0.001 | *** |
|  | Sex | 0.25 | 4.8 | < 0.001 | *** |
| Stay after (pot.) switch (9B, 2<sup>nd</sup>-to-right) | PDS | 0.48 | 3.5 | < 0.001 | *** |
|  | Sex | 0.76 | 2.9 | 0.0036 | ** |
| Asymptotic performance (9B, right) | PDS | 0.25 | 4.2 | < 0.001 | *** |
|  | Sex | 0.098 | 0.9 | 0.39 |  |
| Overall accuracy (9C, left) | Test. | < 0.0001 | 1.2 | 0.24 |  |
|  | Sex | 0.032 | 0.69 | 0.49 |  |
| Response times (9C, 2<sup>nd</sup>-to-left) | Test. | -0.0034 | -5.1 | < 0.001 | *** |
|  | Sex | 0.010 | 1.9 | 0.049 | * |
| Stay after (pot.) switch (9C, 2<sup>nd</sup>-to-right) | Test. | 0.012 | 3.5 | < 0.001 | *** |
|  | Sex | 0.27 | 1.0 | 0.29 |  |
| Asymptotic performance (9C, right) | Test. | 0.0034 | 2.2 | 0.029 | * |
|  | Sex | 0.12 | 1.0 | 0.34 |  |

**Table 11:** Parameter estimates and statistics from hierarchical model fitting, for pubertal predictors (PDS questionnaire, salivary testosterone), for participants under the age of 18. Significance tests against 0 for parameters whose range includes 0, NA otherwise.

| Model | Parameter | $\mu + -sd$ | 95% CI | p-value | sig. |
| --- | --- | --- | --- | --- | --- |
| **PDS** |  |  |  |  |  |
| 4-param. BI | $p_{int}$ | $0.11 + -0.013$ | [0.082, 0.13] | $< 0.001$ | *** |
| | $p_{sd}$ | $0.089 + -0.0085$ | [0.073, 0.11] | 0 | NA |
| | $p_{lin}$ | $0.022 + -0.0096$ | [0.0039, 0.041] | 0.0086 | ** |
| | $\beta_{int}$ | $3.81 + -0.26$ | [3.31, 4.34] | 0 | NA |
| | $\beta_{sd}$ | $1.25 + -0.14$ | [0.98, 1.53] | 0 | NA |
| | $\beta_{lin}$ | $0.31 + -0.16$ | [-0.018, 0.62] | 0.028 | * |
| | $Preward_{int}$ | $0.88 + -0.019$ | [0.84, 0.92] | 0 | NA |
| | $Preward_{sd}$ | $0.060 + -0.011$ | [0.038, 0.082] | 0 | NA |
| | $Preward_{lin}$ | $< 0.001 + -0.010$ | [-0.019, 0.020] | 0.48 | - |
| | $Pswitch_{int}$ | $0.16 + -0.016$ | [0.13, 0.20] | 0 | NA |
| | $Pswitch_{sd}$ | $0.067 + -0.0070$ | [0.053, 0.080] | 0 | NA |
| | $Pswitch_{lin}$ | $-0.0098 + -0.0099$ | [-0.029, 0.0099] | 0.16 | - |
| 4-param. RL | $p_{int}$ | $0.25 + -0.026$ | [0.20, 0.30] | $< 0.001$ | *** |
| | $p_{sd}$ | $0.24 + -0.019$ | [0.20, 0.28] | 0 | NA |
| | $p_{lin}$ | $0.039 + -0.024$ | [-0.0093, 0.087] | 0.054 | - |
| | $\beta_{int}$ | $3.15 + -0.13$ | [2.90, 3.41] | 0 | NA |
| | $\beta_{sd}$ | $1.37 + -0.13$ | [1.12, 1.62] | 0 | NA |
| | $\beta_{lin}$ | $0.41 + -0.13$ | [0.17, 0.66] | $< 0.001$ | *** |
| | $\alpha_{- int}$ | $0.60 + -0.016$ | [0.56, 0.62] | 0 | NA |
| | $\alpha_{- sd}$ | $0.16 + -0.013$ | [0.14, 0.18] | 0 | NA |
| | $\alpha_{- lin}$ | $-0.0155 + -0.017$ | [-0.048, 0.019] | 0.18 | - |
| | $\alpha_{+ int}$ | $0.66 + -0.028$ | [0.61, 0.72] | 0 | NA |
| | $\alpha_{+ sd}$ | $0.35 + -0.034$ | [0.023, 0.15] | 0 | NA |
| | $\alpha_{+ lin}$ | $0.0085 + -0.027$ | [-0.048, 0.059] | 0.38 | - |
| **Testosterone** |  |  |  |  |  |
| 4-param. BI | $p_{int}$ | $0.11 + -0.013$ | [0.081, 0.13] | $< 0.001$ | *** |
| | $p_{sd}$ | $0.089 + -0.0084$ | [0.073, 0.11] | 0 | NA |
| | $p_{lin}$ | $0.02 + -0.010$ | [0.0023, 0.040] | 0.015 | * |
| | $\beta_{int}$ | $3.78 + -0.26$ | [3.29, 4.31] | 0 | NA |
| | $\beta_{sd}$ | $1.28 + -0.14$ | [1.00, 1.55] | 0 | NA |
| | $\beta_{lin}$ | $0.12 + -0.17$ | [-0.20, 0.45] | 0.22 | - |
| | $Preward_{int}$ | $0.88 + -0.019$ | [0.85, 0.92] | 0 | NA |
| | $Preward_{sd}$ | $0.056 + -0.011$ | [0.035, 0.077] | 0 | NA |
| | $Preward_{lin}$ | $-0.0135 + -0.010$ | [-0.033, 0.0081] | 0.90 | - |
| | $Pswitch_{int}$ | $0.16 + -0.016$ | [0.13, 0.19] | 0 | NA |
| | $Pswitch_{sd}$ | $0.067 + -0.0069$ | [0.054, 0.081] | 0 | NA |
| | $Pswitch_{lin}$ | $-0.0082 + -0.010$ | [-0.029, 0.012] | 0.22 | - |
| 4-param. RL | $p_{int}$ | $0.24 + -0.025$ | [0.20, 0.29] | $< 0.001$ | *** |
| | $p_{sd}$ | $0.24 + -0.0195$ | [0.20, 0.28] | 0 | NA |
| | $p_{lin}$ | $0.038 + -0.025$ | [-0.0091, 0.190] | 0.066 | - |
| | $\beta_{int}$ | $3.16 + -0.14$ | [2.89, 3.43] | 0 | NA |
| | $\beta_{sd}$ | $1.42 + -0.13$ | [1.17, 1.69] | 0 | NA |
| | $\beta_{lin}$ | $0.28 + -0.13$ | [0.037, 0.54] | 0.013 | * |
| | $\alpha_{- int}$ | $0.60 + -0.017$ | [0.55, 0.62] | 0 | NA |
| | $\alpha_{- sd}$ | $0.16 + -0.013$ | [0.13, 0.18] | 0 | NA |
| | $\alpha_{- lin}$ | $-0.035 + -0.018$ | [-0.070, -0.0016] | 0.24 | - |
| | $\alpha_{+ int}$ | $0.66 + -0.028$ | [0.61, 0.72] | 0 | NA |
| | $\alpha_{+ sd}$ | $0.10 + -0.030$ | [0.045, 0.16] | 0 | NA |
| | $\alpha_{+ lin}$ | $-0.017 + -0.026$ | [-0.066, 0.036] | 0.015 | * |

#### 6.3.4. Qualitative Model Fit of RL and BI

To test the qualitative fit of our models, we simulated behavior using fitted parameters (from the age-free model; section 4.5.3) and checked whether the simulated behavior was able to reproduce the patterns of interest in the human data (Blohm et al., 2020; Palminteri et al., 2017; Wilson and Collins, 2019). We found that RL and BI models replicated human behavior and age differences, including linear increase in staying after positive outcomes (“+ +” and “-+”), and the inverse-U shape on potential switch trials (red arrow; “+ -” condition). Qualitative (non-significant) sex differences were also captured (suppl. Fig. 15B). Both models also captured quicker switching on switch trials in younger (light green) compared to older participants (blue and grey), and best performance on asymptotic trials in adolescents (greenblue; suppl. Fig. 15A). In summary, both the winning RL and BI model captured human learning curves, as well as sex and age differences, very closely. Simpler, non-winning models, on the other hand, failed to capture human characteristics (suppl. Fig. 17, 16).

**Figure 15:**
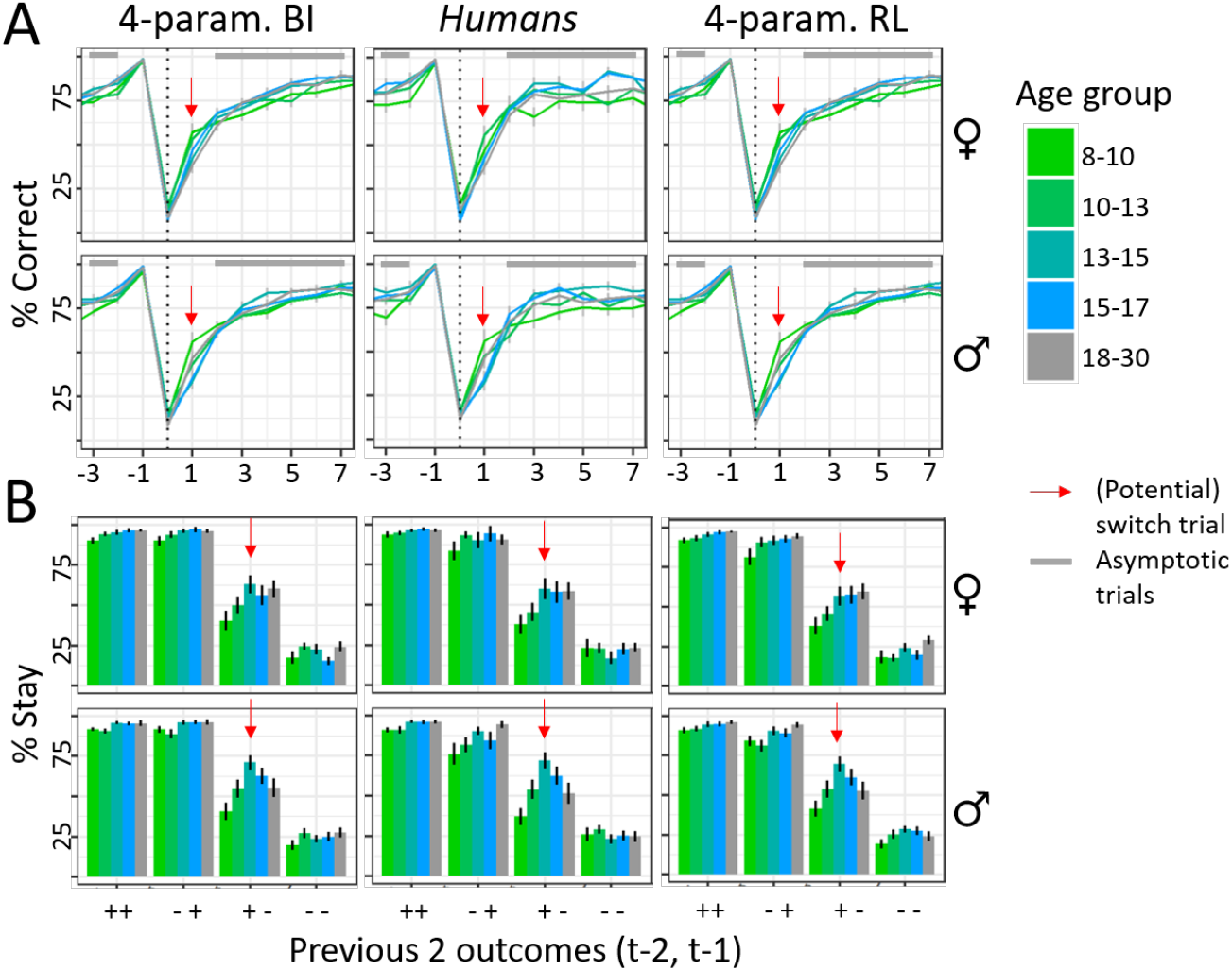
A) Behavior in response to switch trials. Colors refer to age groups, red arrows show switch trials, grey bars trials of asymptotic performance. B) Stay probability in response to outcomes 1 and 2 trials back.

To asses effects of age groups, we tested differences in posterior samples of the age-free model. Statistics are shown in suppl. Table 12.

**Table 12:** Parameter differences between specific age groups. p-values were obtained by assessing means for each parameter for three age groups (8-10, 13-15, and 18-30) and show in how many MCMC samples the group mean of 8-10 year olds (18-30 year olds) was smaller than the group mean of mid- to late adolescence.

| Parameter | Compared groups | p-value | sig. |
| --- | --- | --- | --- |
| $\alpha_-$ | 8-10 vs 13-15 | 0 | *** |
|  | 13-15 vs 18-30 | 0.0045 | ** |
| $p_{reward}$ | 8-10 vs 13-15 | 0.019 | * |
|  | 13-15 vs 18-30 | 0.078 | , |
| $p_{switch}$ | 8-10 vs 13-15 | 0.023 | * |
|  | 13-15 vs 18-30 | 0.13 |  |

**Table 13:** Parameter estimates and statistics from hierarchical model fitting. Significance tests against 0 for parameters whose ranges include 0, NA otherwise.

| Model | Parameter | $\mu + -sd$ | 95% CI | p-value | sig. |
| --- | --- | --- | --- | --- | --- |
| **4-param. RL** | $p_{int}$ | $0.34 + -0.027$ | [0.29, 0.39] | $< 0.001$ | *** |
| | $p_{sd}$ | $0.24 + -0.015$ | [0.21, 0.26] | 0 | NA |
| | $p_{lin}$ | $0.11 + -0.020$ | [0.075, 0.15] | $< 0.01$ | ** |
| | $p_{qua}$ | $-0.050 + -0.020$ | [-0.089, -0.012] | 0.0051 | ** |
| | $\beta_{int}$ | $3.48 + -0.15$ | [3.18, 3.79] | 0 | NA |
| | $\beta_{sd}$ | $1.48 + -0.10$ | [1.29, 1.69] | 0 | NA |
| | $\beta_{lin}$ | $0.36 + -0.11$ | [0.14, 0.57] | $< 0.001$ | *** |
| | $\beta_{qua}$ | $-0.22 + -0.11$ | [-0.42, -0.015] | 0.020 | * |
| | $\alpha_{- int}$ | $0.60 + -0.018$ | [0.56, 0.63] | 0 | NA |
| | $\alpha_{- sd}$ | $0.16 + -0.0093$ | [0.14, 0.18] | 0 | NA |
| | $\alpha_{- lin}$ | $0.011 + -0.015$ | [-0.017, 0.040] | 0.77 | |
| | $\alpha_{- qua}$ | $0.013 + -0.014$ | [-0.013, 0.040] | 0.84 | |
| | $\alpha_{+ int}$ | $0.73 + -0.034$ | [0.66, 0.79] | 0 | NA |
| | $\alpha_{+ sd}$ | $0.081 + -0.021$ | [0.042, 0.12] | 0 | NA |
| | $\alpha_{+ lin}$ | $0.055 + -0.024$ | [0.0045, 0.10] | 0.015 | * |
| | $\alpha_{+ qua}$ | $-0.015 + -0.021$ | [-0.055, 0.027] | 0.25 | |
| **4-param. BI** | $p_{int}$ | $0.13 + -0.013$ | [0.11, 0.16] | $< 0.001$ | *** |
| | $p_{sd}$ | $0.081 + -0.0061$ | [0.069, 0.093] | 0 | NA |
| | $p_{lin}$ | $0.04 + -0.008$ | [0.023, 0.054] | $< 0.001$ | *** |
| | $p_{qua}$ | $-0.02 + -0.007$ | [-0.038, -0.010] | $< 0.001$ | *** |
| | $\beta_{int}$ | $4.27 + -0.27$ | [3.76, 4.83] | 0 | NA |
| | $\beta_{sd}$ | $1.39 + -0.12$ | [1.16, 1.64] | 0 | NA |
| | $\beta_{lin}$ | $0.39 + -0.17$ | [0.054, 0.72] | 0.011 | * |
| | $\beta_{qua}$ | $< 0.001 + -0.16$ | [-0.32, 0.30] | 0.49 | |
| | $p_{reward int}$ | $0.87 + -0.016$ | [0.84, 0.91] | 0 | NA |
| | $p_{reward sd}$ | $0.064 + -0.0087$ | [0.046, 0.081] | 0 | NA |
| | $p_{reward lin}$ | $0.0045 + -0.0096$ | [-0.014, 0.024] | 0.68 | |
| | $p_{reward qua}$ | $-0.0017 + -0.0085$ | [-0.018, 0.015] | 0.43 | |
| | $p_{switch int}$ | $0.16 + -0.014$ | [0.14, 0.19] | 0 | NA |
| | $p_{switch sd}$ | $0.071 + -0.0053$ | [0.062, 0.083] | 0 | NA |
| | $p_{switch lin}$ | $-0.0066 + -0.0095$ | [-0.025, 0.012] | 0.24 | |
| | $p_{switch qua}$ | $0.014 + -0.0082$ | [-0.0013, 0.030] | 0.042 | * |

To evaluate continuous age effects in a statistically sound way, we used a hierarchical Bayesian model that explicitly modeled age effects (the “age-based” model; Fig. 3B). Significant effects (suppl. Table 13) are shown as lines in suppl. Figures 17 and 16.

**Figure 16:**
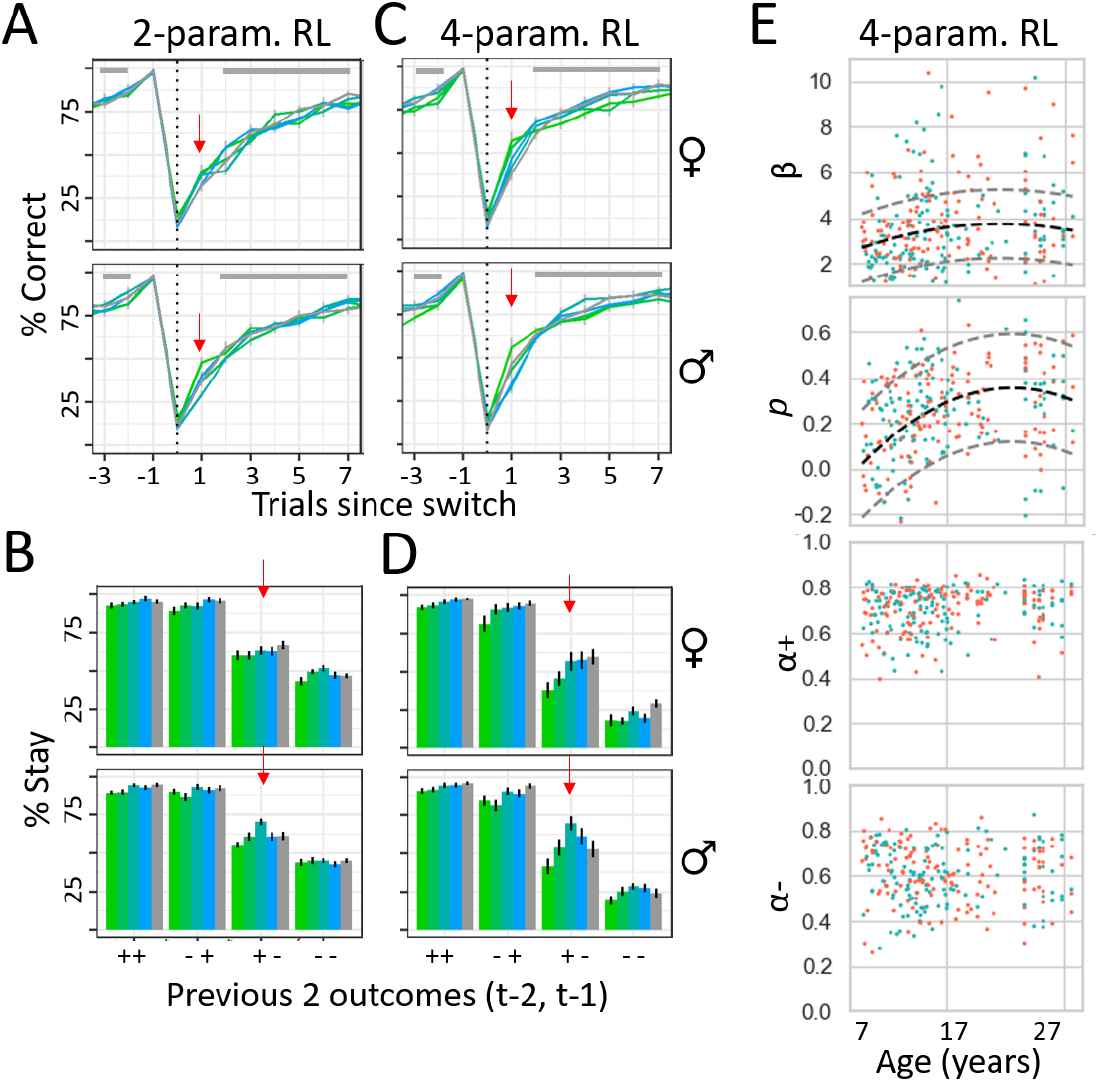
Qualitative fit of different versions of the RL model. Model behavior is shown in the same way as human behavior in suppl. Fig. 15. A-B) Behavior of simulations from the basic, 2-parameter version, with free parameters *α* and *β*. Lacking counterfactual updating and the ability to differentiate positive and negative outcomes, the model was unable to capture the shape of human learning curves and age differences. Colors denote age groups, red arrow (potential) switch trials, and grey bars asymptotic trials, as in suppl. Fig. 15. C-D) Behavior of simulations from the winning, 4-parameter RL model, in which free parameters *β*, *p*, *α*_+_, and *α_−_* were fitted to participants using hierarchical Bayesian model fitting (age-less model; see section 4.5.3). E) Fitted parameters of each individual. Dashed lines show significant age differences (Table 13). This is the same data as summarized in Fig. 4A-D.

**Figure 17:**
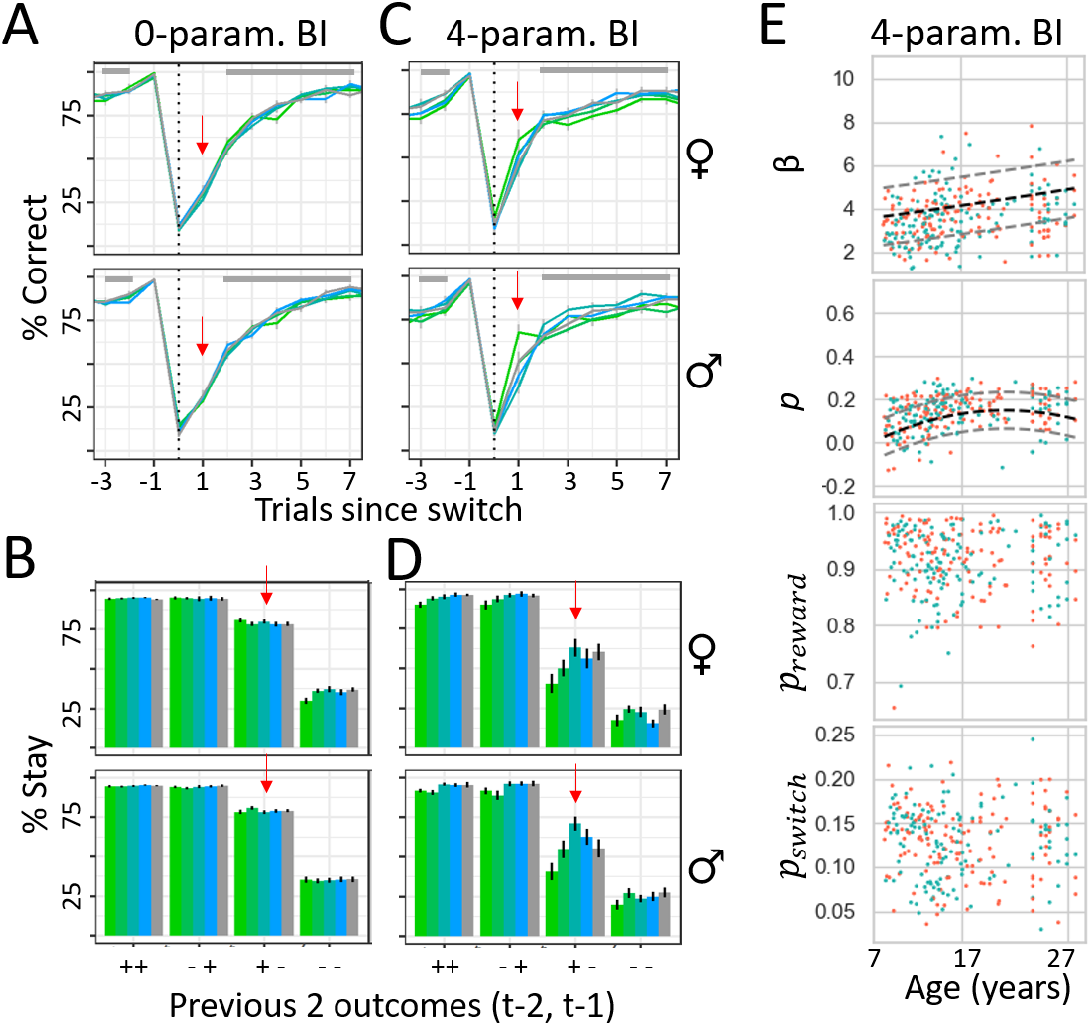
Qualitative fit of different versions of the BI model. Model behavior is shown in the same way as human behavior in suppl. Fig. 15. A-B) Behavior of simulations from the basic, 0-parameter version, in which truthfully *p_reward_* = 0.75 and *p_switch_* = 0.05. Lacking free parameters, the model predicted the same behavior for all participants, being unable to capture age differences. C-D) Behavior of simulations from the winning, 4-parameter version of the BI model, in which free parameters *β*, *p*, *p_reward_*, and *p_switch_* were fitted to participants using hierarchical Bayesian model fitting. To avoid double-dipping into age differences when visualizing the model, we fitted the model *without* access to participants’ age (Methods). E) Fitted parameters of each individual, based on the same model. Dashed lines show age differences when significant (suppl. Table 13). This is the same data as summarized in Fig. 4.

#### 6.3.5. Generate and Recover Model Parameters (Fig. 5A)

In order to assess whether the RL and BI models made the same or different behavioral predictions, we conducted a generate-and-recover test (section 2.3): Artificial behavior is simulated from both models, and the simulated datasets are fitted using both models. Specifically, we simulated one dataset per participant from each model (RL and BI), using the model parameters fitted for the participant (age-free model). We then fitted the simulated data with the RL and BI model (age-free model). We finally calculated WAIC scores and standard errors using PyMC3 (Salvatier et al., 2016). If both datasets are fitted equally well by both models, they are not distinguishable—the behavior they each produce is so similar that both models capture it equally well. If one model fits both behavioral datasets better, it is more appropriate and subsumes the other. If, however, each model fits the artificial dataset better that was generated by its own class (e.g., *RL*↔*RL*), both models must produce different behaviors to explain why the corresponding model captures it more neatly. This pattern was the case for our models: Based on human-fitted parameter values for simulation (Heathcote et al., 2015; Wilson and Collins, 2019), each model fit its own simulated dataset better than to the other model’s (Fig. 5A). This confirms that the winning RL and BI models were distinguishable, i.e., predicted different behaviors.

Comparable results were obtained when using the more classical generate-and-recover method of assessing the number of best-fitted models based on maximum likelihood (suppl. Fig. 18), rather than hierarchical Bayesian model fit (WAIC; Fig. 5A).

**Figure 18:**
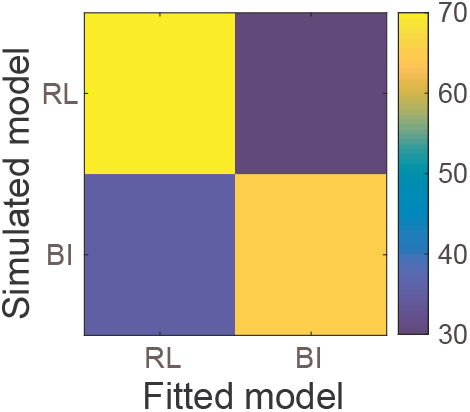
Number of simulated datasets (out of 100) of each model (y-axis) that were best fit using each of the two models (RL and BI; x-axis). Lighter colors indicate larger fractions and highlight the diagonal of the confusion matrix, showing that RL simulations were best recovered by the RL model and BI simulations by the BI model, using maximum likelihood.

#### 6.3.6. Trial Types of Behavioral Difference for RL versus BI

Further analyses showed that the differences between RL and BI could be traced to specific types of decision situations: The RL model is more likely to stay with a choice after receiving two consecutive rewards than after receiving just a single reward because the action value is increased twice in the former case, but only once in the latter. To assess this, we used a t-test to compare the probability of RL model simulations (based on human-fitted parameters) to stay between both cases (*t* = 2.6, *p* = 0.010). The BI model, however, is equally likely to stay in both cases (*t* = −0.5, *p* = 0.6) because a single reward already leads to maximally certain state inference, and another reward cannot increase this probability further. This analysis provides a concrete example of how the RL and BI models differ in terms of behavior, confirming that they do not make identical predictions.

#### 6.3.7. Between-Model Parameter Similarities, Assessed Using Regression (Fig. 5C)

The correlation analysis in section 2.4 showed that both models captured similar processes using different *individual* parameters, but similar processes might also be captured in the interplay between *several* parameters. To investigate this possibility, we used linear regression to evaluate how well we could predict each parameter based on the parameters and one-way parameter interactions of the other model. This analysis revealed that 7 of 8 parameters could be predicted almost perfectly (Fig. 5C), showing that the interplay between parameters in one model captured almost all variance in almost every parameter in the opposite model. In other words, fitting the RL model to participants’ data allowed us to nearly perfectly predict participants’ BI parameters, without fitting the BI model. Parameter *α*_+_ (RL) was again an exception, with only small amounts of variance captured by BI parameters, suggesting that it reflected mechanisms that were unique to the RL model. These mechanisms might increase the versatility of the RL model, and possibly account for the slightly better numerical fit of the RL model to human (Table 2) and simulated data (Fig. 5A). In sum, in addition to significant similarities between individual parameters, the RL and BI models showed even greater similarities in terms of cognitive processes that were captured in the interactions between multiple parameters. This suggests that both models captured very similar cognitive processes, albeit without reaching identity (e.g., parameter *α*_+_).

We ran eight different regression models, predicting each parameter from the 4 parameters of the opposite model, as well as their one-way interactions, using linear regression in R (RCoreTeam, 2016). Fig. 5C shows the explained variance (*R*^2^) of each model.

#### 6.3.8. Details on the PCA Analysis

We conducted a PCA on the joint parameter space of our winning RL and BI models in the hope of identifying model-general factors (PCs) that explain age differences in cognitive processing. The crucial step in this analysis is to interpret the resulting PCs. PCs are often interpreted through the weights (factor loadings) that each raw feature (model parameter) has on the PC (a PC is just a linear combination of raw features). In our case, this approach was impeded by the fact that model parameters are themselves difficult to interpret because their roles are influenced by many factors, including the underlying task (Eckstein, Master, et al., 2021) and computational model (Sugawara and Katahira, 2021), which makes them less suitable to anchor the meaning of PCs.

For this reason, we devised the following simulation approach: We simulated data from our computational models based on the obtained principal components (PCs) in order to visualize the role of each PC. It is common practice to simulate data based on small or large values of a parameter (e.g., smaller or larger decision noise *β*) to assess the role of this parameter for model behavior (e.g., better or worse performance). We similarly simulated data based on smaller or larger values of each PC to clarify the precise role of each PC: We calculated two sets of parameters for each PC, one that represented high levels of this PC (“plus”), and one that represented low values (“minus”). Low levels were determined by subtracting 4 times the inverse-z-scored factor loading of a PC (center) from the population mean of each parameter; low levels were determined by adding it. Suppl. Table 14 shows these two sets of parameters. (For PC2 of the BI model, we added and subtracted 2 times the factor loading instead, to ensure *p_reward_ <* 1.) We then simulated behavior based on the resulting parameters to assess the effect of low versus high values of each PC.

**Table 14:** Parameters used in suppl. Fig. 19 to visualize the role of PCs.

| | $p$ (RL) | $\beta$ (RL) | $\alpha_-$ | $\alpha_+$ | $p$ (BI) | $\beta$ (BI) | $p_{reward}$ | $p_{switch}$ |
| --- | --- | --- | --- | --- | --- | --- | --- | --- |
| PC1 (plus) | 0.57 | 6.95 | 0.45 | 0.87 | 0.26 | 5.67 | 0.84 | 0.07 |
| PC1 (minus) | 0.04 | 0.10 | 0.80 | 0.65 | 0.02 | 1.72 | 0.98 | 0.20 |
| PC2 (plus) | 0.06 | 2.65 | 0.31 | 0.64 | 0.10 | 2.98 | 0.84 | 0.12 |
| PC2 (minus) | 0.54 | 4.41 | 0.94 | 0.89 | 0.18 | 4.41 | 0.98 | 0.15 |
| PC3 (plus) | 0.76 | 0.49 | 0.57 | 0.74 | 0.29 | 1.87 | 0.85 | 0.19 |
| PC3 (minus) | -0.16 | 6.56 | 0.68 | 0.78 | -0.01 | 5.52 | 0.97 | 0.08 |
| PC4 (plus) | 0.15 | 1.68 | 0.58 | 1.19 | 0.10 | 3.06 | 0.88 | 0.14 |
| PC4 (minus) | 0.45 | 5.38 | 0.67 | 0.33 | 0.18 | 4.33 | 0.94 | 0.13 |
| Parameter mean | 0.30 | 3.53 | 0.62 | 0.76 | 0.14 | 3.69 | 0.91 | 0.13 |

This analysis revealed that PC1, capturing the largest proportion of parameter variance, reflected a broad measure of behavioral quality: Low values of PC1 led to low performance and lacked differentiation between different outcome histories, while high values led to high performance and efficient responses that were in tune with outcome histories (suppl. Fig. 19A; suppl. Table 14). PC1 factor loadings revealed that low behavioral quality was related to larger-than-average values of *α_−_* (RL), which likely led to premature switching due to the over-sensitivity to recent negative outcomes. Low behavioral quality was also due to larger-than-average values of *p_reward_* and *p_switch_* (BI), which created overly deterministic and overly volatile mental models of the task; whereas an overly deterministic task model leads to pre-mature switching after negative outcomes (because negative outcomes only arise in deterministic tasks when contingencies have switched), and an overly volatile task model leads to a reduced reliance on past outcomes (because frequent task switches mean that past information is soon outdated; suppl. Fig. 19A, center). High behavioral quality, on the other hand, was caused by larger-than-average values of *α*_+_ (RL), which underlies the quick learning from positive outcomes, and therefore reliable staying behavior after (diagnostic!) outcomes. High behavioral quality was also caused by larger-than-average values of *p* (RL and BI), which increased choice persistence, facilitating repetition of non-rewarded actions; and of larger-than-average values of *β* (RL and BI), which reduced decision noise, allowing for a more direct translation of beliefs (BI) or action values (RL) into choices.

PC2 represented integration time scales: Low values of PC2 (short time scales) led to win-stay behavior—defined as immediate switching after negative outcomes and consistent staying after positive outcomes—, which resulted in poor performance on asymptotic trials (suppl. Fig. 19B, left). High values of PC2, on the other hand, led to increasingly slow behavioral switches, resulting in poor performance on switch trials (suppl. Fig. 19B, right). In order to achieve high performance on both asymptotic and switch trials, participants needed to find the appropriate balance between both ends on this spectrum. PC3 captured responsiveness to task outcomes: Low values of PC3 led to a lack of differentiation between outcome histories and slow behavioral switching (suppl. Fig. 19C, right), whereas high values led to extremely consistent win-stay-lose-shift behavior (suppl. Fig. 19C, left). PC4 uniquely captured RL parameter *α*_+_, i.e., the tension between slow versus fast updates when integrating positive outcomes (suppl. Fig. 19D). Suppl. Figures 19B, C, and D (center) show which model parameters drove the behavior of PC2-4.

**Figure 19:**
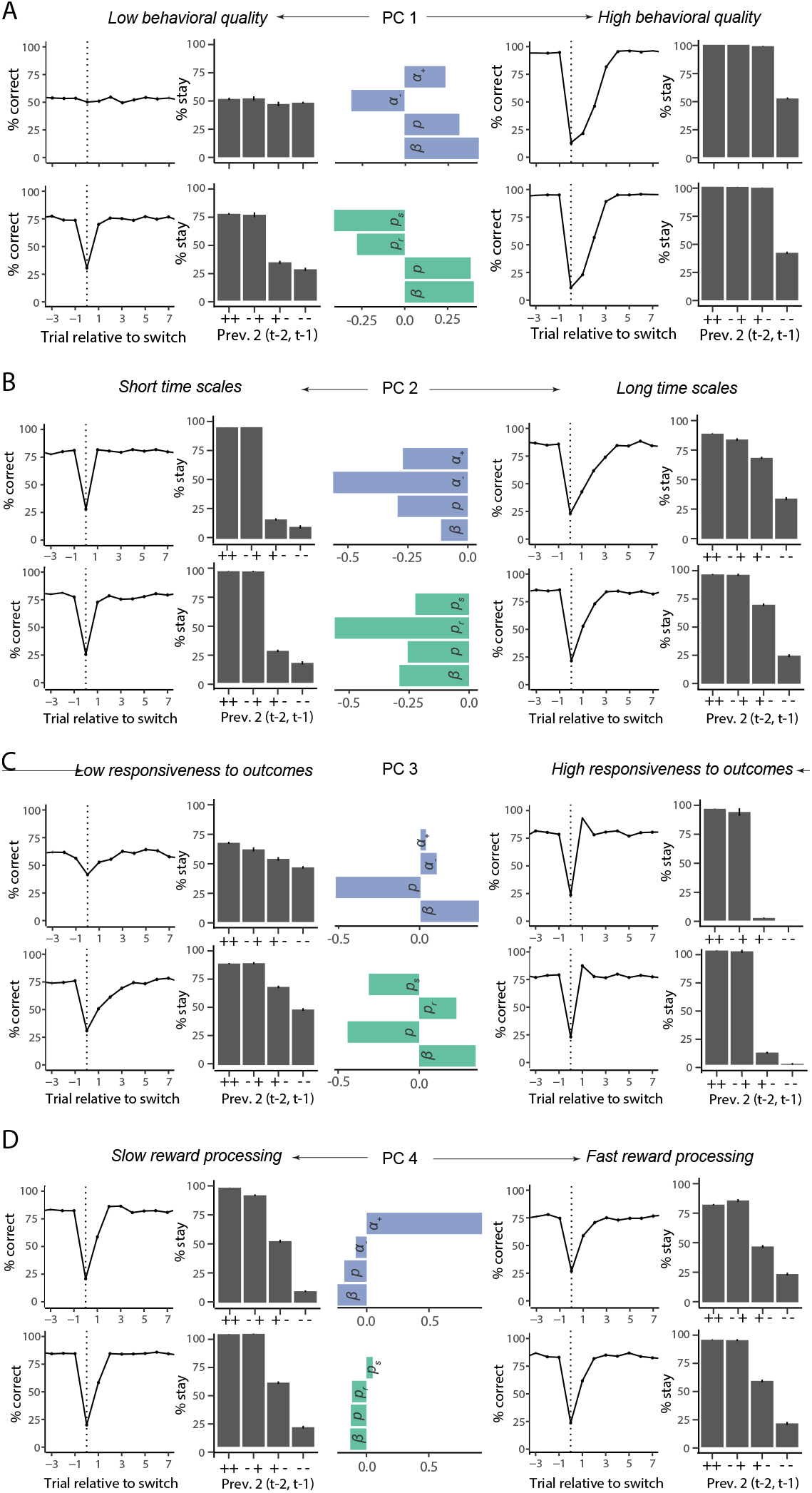
Determining the role of each PC for behavior. The figure shows simulated behavior based on low (left) and high (right) values of each PC. Parts A-D) show the results for PCs1-4.

To address the main question of our study, we also assessed age differences in PCs. Table 15 shows the results of this analysis.

**Table 15:** Results of t-tests on PC2 and PC4. df: Welch-adjusted degrees of freedom.

| Comparison | $t$ | df | $p$ | Sig. |
| --- | --- | --- | --- | --- |
| PC2 (8-15 vs. 15-30) | 3.44 | 266.2 | < 0.001 | *** |
| PC4 (8-13 vs. 13-17) | 2.28 | 176.8 | 0.047 | * |
| PC4 (13-17 vs. 18-30) | 2.49 | 176.6 | 0.028 | * |

### 6.4. Supplemental Discussion

#### 6.4.1. Potential Effect of Recruitment on Results

As explained in the Discussion, it is not impossible that recruitment differences affected our results. However, speaking against this possibility, recruitment processes were identical for children, adolescents, and community adults, limiting the possibility that the observed age differences were due to recruitment. The only age group that was recruited differently were college students (see section 4.1); however, given the competitive nature of the college, we would expect college students to perform *better* than adolescents and not worse. Furthermore, removing college students does not affect the observed behavioral peak in adolescence. Lastly, the adolescent peak was specific to the current task, and did not arise in two structurally-similar learning tasks participants performed in the same session (Master et al., 2020; Xia et al., 2020; for side-by-side comparison, see Eckstein, Master, et al., 2021; Eckstein, Wilbrecht, et al., 2021). Both other tasks lacked the reversal aspect, suggesting that adolescents are specifically adapted to reversal, in accordance with the similarity in findings in van der Schaaf et al., 2011, a deterministic reversal task.

#### 6.4.2. Different Models at Different Ages?

Previous studies have shown that participants of different ages sometimes are better fitted by different computational models, suggesting that they might employ different cognitive mechanisms (e.g., Palminteri et al., 2016). Could the same apply to our study? For example, previous studies have reported age-based increases in “model-based” (Decker et al., 2016) and counterfactual learning (Palminteri et al., 2016), which might reflect an improved mental task model. Accordingly, one might expect that in our study, children’s cognitive processes would resemble a simple incremental RL model, whereas adolescents’ would resemble the mental-model-based— and more optimal—BI model. Even though this is a justified question, it is unlikely that different models applied to different age groups in our study, given that both models captured the behavior of all age groups equally well in model validation. Compared to previous studies that showed age differences in model types, the greater flexibility of our models in terms of the number of free parameters and augmentations might have allowed them to capture more age differences, obliterating the need to change the model itself.

## Notes

### Competing Interest Statement

The authors have declared no competing interest.

### Summary of Updates

This version of the manuscript has been revised to emphasize results showing the adolescent peak in behavior. The article has also been shortened.

